# Trajectories of brain volume change over 13 years in chronic schizophrenia

**DOI:** 10.1101/2019.12.17.879429

**Authors:** Claudia Barth, Kjetil N. Jørgensen, Laura A. Wortinger, Stener Nerland, Erik G. Jönsson, Ingrid Agartz

## Abstract

**Importance:** Schizophrenia is a leading cause of disability worldwide, with an illness course that putatively deteriorates over time. Whether the notion of a progressive brain disease holds in its chronic stage is debated.

**Objective:** To investigate brain volume change and the impact of iatrogenic factors in chronic schizophrenia patients (duration of illness at baseline 16.17 ± 8.14 years) and controls over 13 years.

**Design:** Participants were recruited as part of the Human Brain Informatics study. Data acquisition took place between 1999 and 2018, including baseline, 5- and 13-years follow-up.

**Setting:** Naturalistic longitudinal case-control study.

**Participants:** The sample consisted of 143 participants, of whom 64 were patients with chronic schizophrenia (20% female, mean age at baseline 40.5 ± 7.7 years) and 79 healthy controls (37% female, mean age at baseline 42.8 ± 8.4 years). T1-weighted structural imaging and information about medication use were obtained at each time point.

**Exposure:** Antipsychotic medication and other prescribed drugs.

**Main Outcome(s) and Measure(s):** Individual total and tissue-specific brain volumes, as well as two-time point percentage brain and ventricle volume change.

**Results:** Patients had lower total brain volume at baseline. Yet, trajectories in total brain volume and gray matter volume loss as well as ventricular enlargement did not differ relative to controls. White matter volume was similar between groups at baseline and 5-year but diverged between 5-year and 13-year follow-up, with accelerated loss in patients. While antipsychotic exposure did not show an association with brain volume loss over time, higher medication load was associated with lower brain volume across time points. Patients on second-generation antipsychotics alone showed lowest total brain volume, only after accounting for add-on drug use.

**Conclusion and Relevance:** We found limited evidence for progressive brain volume loss in chronic schizophrenia, beyond normal aging. Stable differences in patient brain volumes relative to controls may primarily occur during the first years of illness. All prescribed drugs need to be considered when examining the impact of antipsychotic medication on brain structure.

**Key Points:** *Question:* Is chronic schizophrenia associated with progressive brain volume loss beyond normal aging?

*Findings:* While brain volume was lower at baseline, patient trajectories of brain volume change over a 13-year follow-up period differed little from healthy individuals. Small effects indicated greater white matter volume loss in patients during the late phase of follow-up. Stable differences in patient brain volumes seem explicable by antipsychotic medication class and respective add-on drugs.

*Meaning:* We found limited evidence of progressive brain volume loss, beyond normal aging, in chronic schizophrenia over 13 years.

## Introduction

Schizophrenia is a leading cause of disability worldwide, and its ranking as a global burden of disease has not changed in 30 years ^1^. Despite decades of research, treatment remains inadequate, and the illness course putatively deteriorates over time.

Evidence supporting the notion of progressive trajectories stems from multiple studies, including meta-analyses, stating enlargement of the ventricular system and greater gray matter volume reduction of fronto-temporal and limbic regions in schizophrenia patients compared to controls ^2-4^. Studies showing significant brain volume loss span the prodromal state ^5^, first episode psychosis ^6^, first episode schizophrenia ^4^, and chronic schizophrenia ^7^. These studies suggest that progressive tissue loss primarily occurs in the first phases of the illness ^8^. On the contrary, other studies report a lack of longitudinal brain changes in first episode psychosis patients ^9,10^.

Despite mixed results, progressive brain tissue loss seems closely linked to the disease onset. Beyond the early stages of illness, reduction in brain volume seems to stabilize and to be more similar to what can be expected with normal aging ^8^. Results of the Iowa longitudinal study showed that while brain tissue loss in first episode schizophrenia was greatest during the first interscan interval (2 years), changes in brain volume in patients did not differ significantly from those in controls during the second (5-6 years) and third follow-up (8-9 years) ^4^. Another finding by Vita and colleagues was that longer duration of illness was associated with smaller differences in volume change over time between patients and controls ^8^. These findings disagree with the progressive disease perception, at least in its later stages.

The contribution of iatrogenic factors, such as antipsychotic use, to the extent and nature of brain changes in schizophrenia has been extensively discussed. Some authors argue that antipsychotic use may moderate or even cause progressive brain volume loss ^8,11,12^, whereas others state no such relationship ^5^. Emerging evidence suggests that class of antipsychotic drug could be critical: First-generation antipsychotics (FGA), but not second-generation antipsychotics (SGA), may reduce gray matter in different brain regions ^13^. Contrary, SGA has been found to not only reduce, but to partly counteract progressive loss of gray matter tissue, predominantly in the temporal lobe ^8^. However, results are mixed, and a recent systematic review and meta-analysis suggests no clear differences between FGA and SGA exposure and brain volume change ^14^.

Both FGA and SGA have been linked to severe side effects, though with different propensities for the kind of unfavorable impacts. FGA are known to cause a variety of extra-pyramidal symptoms, whereas SGA have been associated with weight gain and metabolic side effects. Add-on drugs are common ^15^ and may relieve side effects. Although antipsychotics emerged in the 1950s with FGA and 1980s with SGA, the long-term consequences of these drugs and their respective add-ons on brain structure are still largely unknown.

Here, using repeated magnetic resonance imaging (MRI) over the course of 13 years, we investigated whether progressive brain volume loss occurs beyond aging in chronic schizophrenia, and how tissue loss is associated with medication use in both long-term-treated patients and healthy controls. We focused on the impact of add-on drugs, classified using the anatomical therapeutic chemical (ATC) classification system, and antipsychotic medication. We hypothesized that patients would show a trajectory of greater brain volume loss compared to healthy participants.

## Methods and Materials

### Participants

Participants were recruited as part of the Human Brain Informatics study at Karolinska Institutet, Stockholm, Sweden. Data acquisition took place between 1999 and 2018, including baseline (T0) and two follow-ups at ∼5-year (T1) and ∼13-year (T2). Patients were recruited from outpatient clinics managed by the Stockholm County health care organization, covering catchment areas within North-Western Stockholm County. Healthy controls were recruited based on population registers and among hospital staff. All participants provided written informed consent to participate. The study was approved by the Research Ethics Committee at Karolinska Institutet.

A flowchart of the study sample is provided in Figure 1. Sample demographics and clinical characteristics at T0 are reported in Table 1. Details about study inclusion and exclusion, as well as information about the study attrition (Table S1), are stated in the supplementary materials.

**Table 1.**
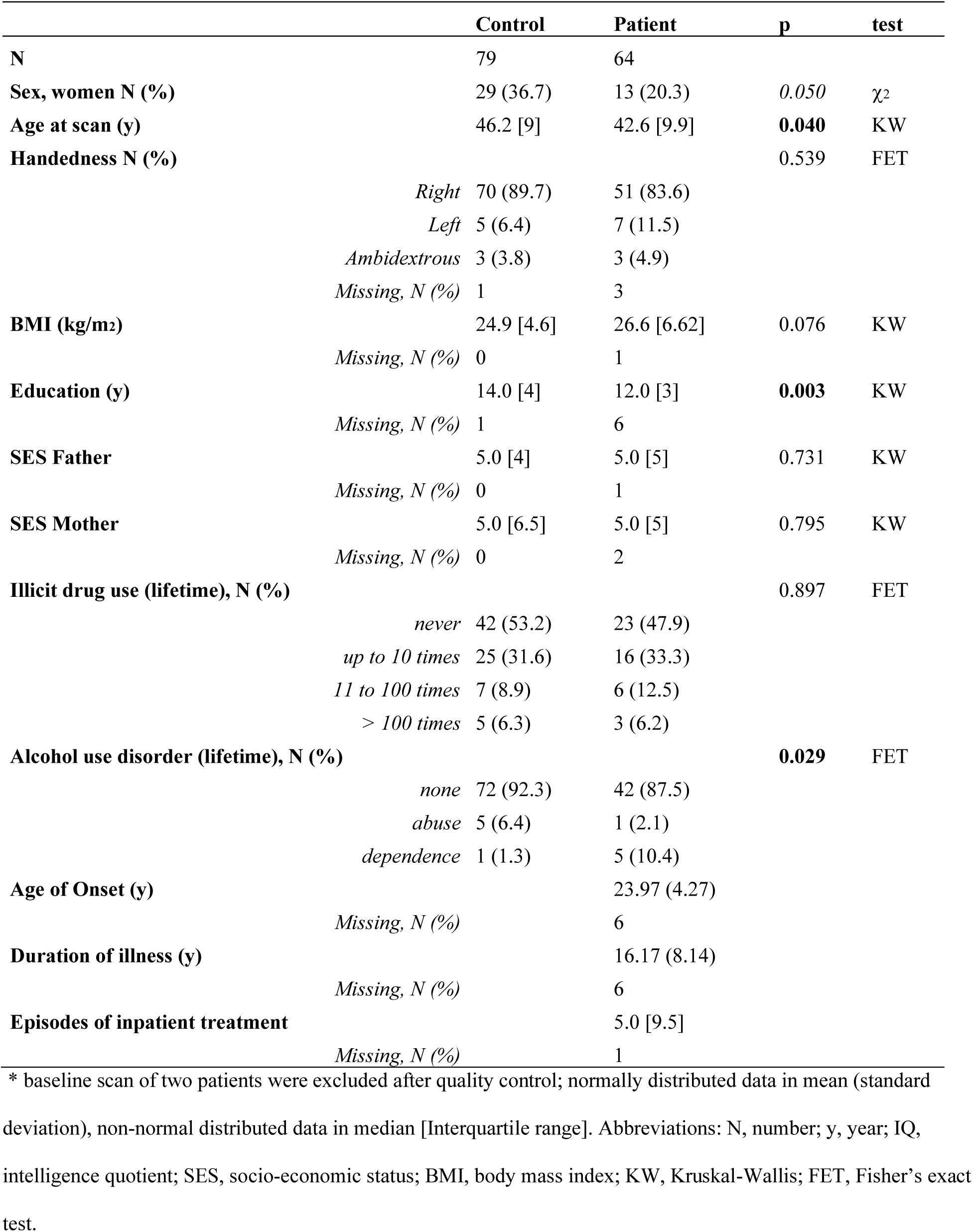
Sample demographic and clinical data at baseline.

**Figure 1.**
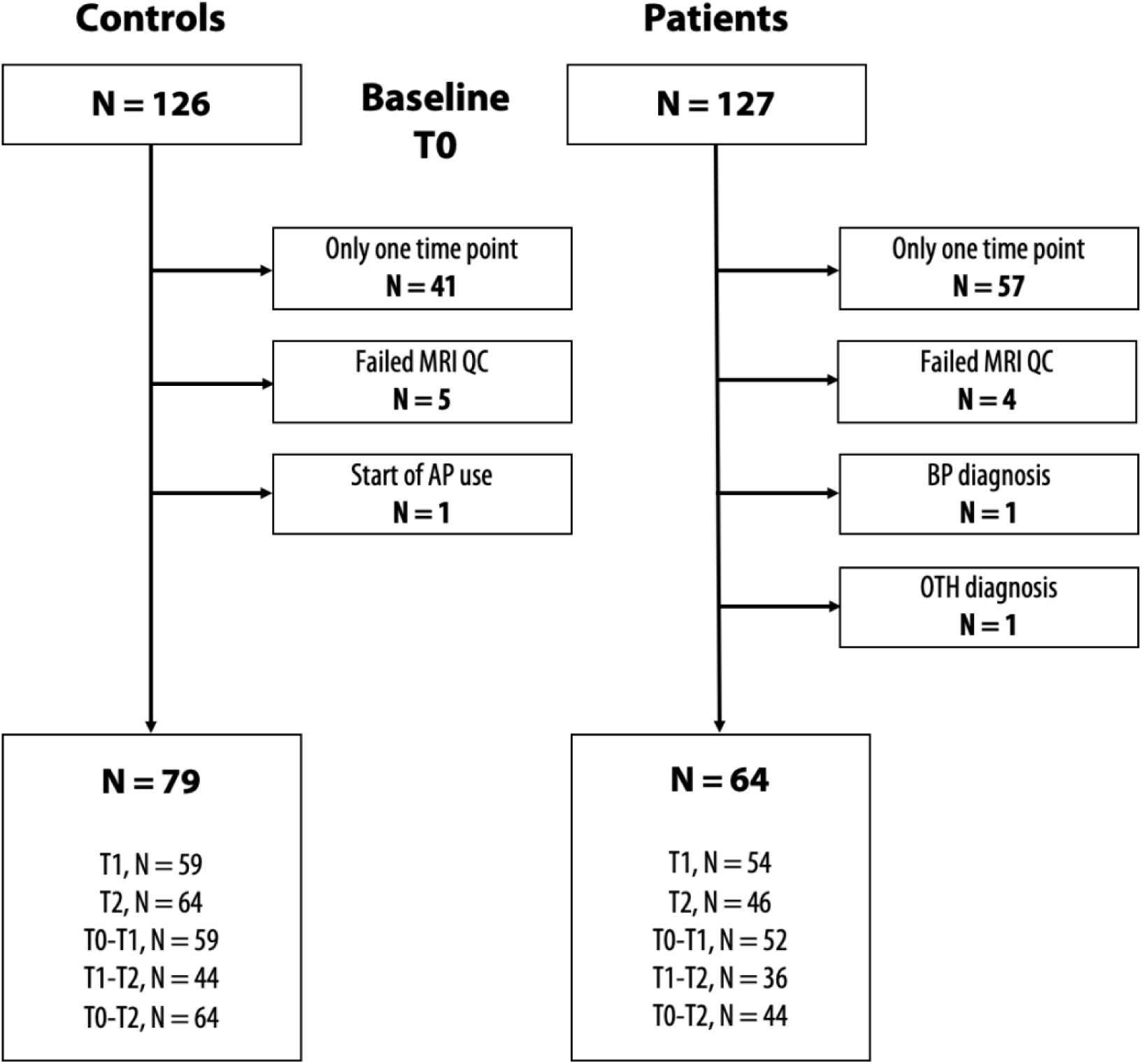
Flowchart of study inclusion and sample size (N) for each time point. The sample size for each time point varies as some participants did not participate either at T1 or T2. A few participants only participated at T0 and T2, not at T1. Abbreviations: T0, baseline; T1, first follow-up; T2, second follow-up; MRI, magnetic resonance imaging; QC, quality control; AP, antipsychotics; BP, bipolar disorder; OTH, opioid induced psychotic disorder, with delusions.

A final sample of 143 participants at T0 (79 controls, 64 patients) was entered into the statistical analysis. Included patients were diagnosed with schizophrenia (n = 49), schizoaffective disorder (n = 11) and psychosis not otherwise specified (n = 4).

### Clinical measures

Symptom rating was evaluated using the Scale for the Assessment of Negative Symptoms (SANS) ^16^ and the Scale for the Assessment of Positive Symptoms (SAPS) ^17^. Global Assessment of Functioning (GAF) was administered for all participants to measure general functioning level^18^. Alcohol and illicit drug use were also evaluated by interview ^19^.

### Medication

For patients, current use of antipsychotic medication was recorded at each time point and converted into chlorpromazine equivalents (CPZ)^20^. The number and generic name of antipsychotics used over time is reported in Table 2. For both patients and controls, use of non-antipsychotic drugs was transformed according to the first level of the ATC classification system^21^ (Table S2, anatomical main group), and shown in depth for the most frequently used ATC main groups using the second level of the ATC Classification System (Table S3, therapeutic subgroup).

**Table 2.** Antipsychotic medication used, and clinical variables stratified by time point in patients.

| **Antipsychotic** | **Class** | **T0, N (%)** | **T1, N (%)** | **T2, N (%)** |
| --- | --- | --- | --- | --- |
| Quetiapine | SGA | 0 (0) | 2 (3.1) | 3 (4.7) |
| Aripiprazole | SGA | 0 (0) | 6 (9.4) | 6 (9.4) |
| Olanzapine | SGA | 14 (21.9) | 14 (21.9) | 13 (20.3) |
| Risperidone | SGA | 6 (9.4) | 5 (7.8) | 3 (4.7) |
| Clozapine | SGA | 7 (10.9) | 7 (10.9) | 8 (12.5) |
| Paliperidone | SGA | 0 (0) | 0 (0) | 3 (4.7) |
| Ziprasidone | SGA | 0 (0) | 0 (0) | 1 (1.6) |
| Haloperidol | FGA | 16 (25) | 13 (20.3) | 10 (15.6) |
| Zuclopenthixol | FGA | 9 (14.1) | 4 (6.2) | 5 (7.8) |
| Perphenazine | FGA | 8 (12.5) | 4 (6.2) | 7 (10.9) |
| Flupentixol | FGA | 1 (1.6) | 2 (3.1) | 2 (3.1) |
| Chlorpromazine | FGA | 1 (1.6) | 0 (0) | 0 (0) |
| Thioridazine | FGA | 1 (1.6) | 0 (0) | 0 (0) |
| No antipsychotics |  | 5 (7.8) | 6 (9.4) | 4 (6.2) |
| Polypharmacy |  | 7 (10.9) | 15 (23.4) | 18 (28.1) |
| Missing |  | 0 (0) | 10 (15.6) | 15 (23.4) |
| **Clinical variables** |  | **T0** | **T1** | **T2** |
| Current CPZ |  | 213.96 [187.84] | 238.01 [189.08] | 303.03 [258.35] |
| Current CPZ, SGA |  | 231.48 [207.32] | 233.64 [105.26] | 319.79 [192.23] |
| Current CPZ, FGA |  | 179.37 [165.14] | 163.04 [180.89] | 163.04 [170.1] |
| SANS, composite score |  | 20.00 [15.50] | 25.50 [18.25] | 28.00 [21.00] |
| *Missing* |  | 25 | 12 | 15 |
| SAPS, composite score |  | 7.00 [10.75] | 8.00 [12.25] | 8.00 [12.00] |
| *Missing* |  | 26 | 12 | 15 |
| GAF |  | 50.00 [20.00] | 50.00 [10.00] | 40.00 [10.00] |
| *Missing* |  | 15 | 11 | 15 |

\*Median [Interquartile range], Abbreviations: SGA, second-generation antipsychotics; FGA, first-generation
antipsychotics; CPZ, chlorpromazine equivalent; N, number; T0, baseline; T1, first follow-up; T2, second follow-up;
SANS, scale for the assessment of negative symptoms; SAPS, scale for the assessment of positive

### MRI data acquisition

MRI data at T0 and T1 was collected using a General Electric 1.5-Tesla Signa HDxt scanner equipped with an 8-channel head coil. At T2, MRI images were acquired at General Electric 3-Tesla Discovery MR750 scanner equipped with a 32-channel head coil. Details see supplementary materials. A trained neuroradiologist evaluated all MRI scans to rule out clinical pathology.

### Structural data processing

T1-weighted imaging data was processed in the FMRIB software library (https://fsl.fmrib.ox.ac.uk/fsl/fslwiki). SIENAX was used to calculate individual total brain volume (TBV) and gray matter, white matter and ventricular cerebrospinal fluid (CSF) volumes at each time point, normalized for head size ^22,23^. To cross-validate SIENAX results, we used SIENA to estimate two-point percentage brain volume change (PBVC) and ventricular volume change (PVVC) from T0 to T2, T0 to T1 and T1 to T2 ^22,23^. For greater regional specificity, we performed analyses of voxel-wise edge displacements. Brain extraction was optimized for both SIENAX and SIENA by pre-masking the T1-weighted images (see supplementary materials for details).

### Statistical analysis

Statistical tests were conducted in R, version 3.5.2.

#### a. Demographic and clinical data

Group differences in demographic and clinical characteristics were assessed using Pearson’s Chi-Squared Test for categorical data, and Student’s t-Tests and Kruskal-Wallis Rank Sum Test for normal and non-normal continuous data, respectively. Changes in clinical scores and medication dosage among patients over time were tested using repeated measures linear mixed-effect model (LMEM) ^24,25^. Fixed factors included the clinical variable of interest, time as continuous variable to account for different interscan intervals, baseline age and sex. As a random factor, intercepts for participants were entered.

To examine the number of 1^st^ level ATC co-medication per participant as a function of group and time, we fitted Bayesian generalized linear mixed-effects models with Poisson distribution^26^. Fixed factors were 1^st^ level ATC code counts and time (continuous) x group interaction. As random effect, we specified intercepts for participants (more details, see supplementary materials).

### Individual brain tissue volume using SIENAX

To assess the relationship between volume measures and group status over time, we fitted separate LMEMs for the dependent variables TBV, gray matter volume, white matter volume and ventricular CSF. As fixed effects, we entered time (continuous) x group interaction, time^2^, baseline age, age^2^, sex, body mass index (BMI) and scanner into the model. Time^2^ and age^2^ were added to model any non-linear effects of time and age on brain volume. As a random effect, we specified intercepts for participants.

#### b. Percentage brain and ventricular volume change using SIENA

Multiple linear regression models were fitted to predict PBVC and PVVC between each time point comparisons, separately. The model specification was similar to the SIENAX LMEM, by entering group status, BMI-change (absolute change), baseline age, age^2^, time (continuous), time^2^ and sex as independent variables.

To obtain a regional measure of significantly atrophic brain edge points between groups, we ran voxel-wise statistics, co-varied for baseline age, time (continuous), sex and BMI change, using a nonparametric permutation-based approach (*randomise*, FSL, 10000 permutations). All variables were mean-centered. The statistical threshold was set at p < 0.05, after family-wise error correction for multiple comparisons using threshold-free cluster enhancement.

#### c. Relationships between clinical variables and SIENAX output

In patients, regression slopes for TBV and SANS, SAPS, GAF and CPZ were assessed with repeated measures correlations ^27^, which determines the overall within-individual relationship among paired measures assessed on multiple occasions ^27^. If a variable of interest showed a significant correlation with TBV, follow-up correlations were performed with these variables and tissue-specific measures.

To examine the effects of antipsychotic classes on TBV, i.e. taking only FGA, only SGA or both, we specified a LMEM with antipsychotic class (categorical), time (continuous), age at baseline, sex, BMI and scanner as fixed factors and intercept for participant as random factor. In addition, as a proxy of cardiovascular- or nervous-system related illnesses, we also included ATC-C and - N drug counts, respectively, as fixed factors in separate models to assess their impact on brain volume measures.

#### d. Relationships between medication load and PBVC

To assess whether CPZ at T0, cumulative CPZ for all time points and CPZ change between time points predicted PBVC in patients, we fitted separate multiple linear regression models using the same specification as the main model (see c).

## Results

### a. Demographic and clinical data

#### Sample demographics and clinical characteristics are reported in Table 1

The attrition sample did not differ significantly from the included sample (see supplementary materials, Table S1). In patients, positive symptoms (SAPS) remained stable over time, while negative symptoms increased (SANS, Estimate=0.35, S.E.=0.16, t=2.15, p=0.04,) and global functioning decreased (GAF, Estimate=-0.47, S.E.=0.12, t=-3.79, p=2.64e-04, see supplementary material Figure S1). Antipsychotic dosage (CPZ) level significantly increased over time (Estimate=0.03, S.E.=7.47e-03, t=3.87, p=2.11e-04). The change in CPZ levels was not driven by significant changes in either FGA or SGA use over time. Yet, combination use of both SGA and FGA together seems to increase over time relative to FGA or SGA use alone (see supplementary materials Figure S2).

Apart from antipsychotic medication, nervous system (ATC-N) and cardiovascular system (ATC-C) medications were the most commonly used drug classes in both patients and controls (see Figure 2). However, ATC-N drugs were more often used in patients than controls (Estimate _Patients_=3.32, S.E.=0.61, t=5.47, p=4.52e-08). In both patients and controls, we found an increase in use over the follow-up periods for ATC-N (Estimate=0.11, S.E.=0.05, t=2.26, p=0.02), ATC-C (Estimate=0.16, S.E.=0.05, t=3.62, p=2.92e-04) and blood and blood forming organs medications (ATC-B; Estimate=0.19, S.E.=0.09, t=2.16, p=0.03). For other drug classes, no main effect of time was found. Prescribed non-antipsychotic nervous system drugs were analgesics, antiepileptics, anti-parkinson drugs, psycholeptics and psychoanaleptics; and prescribed cardiovascular system drugs were diuretics, beta blocking agents, calcium channel blockers, agents acting on the renin-angiotensin system and lipid modifying agents (see supplementary materials, Table S3). Medication use stratified by antipsychotic medication class in patients is displayed in the supplementary material, Figure S3.

**Figure 2.**
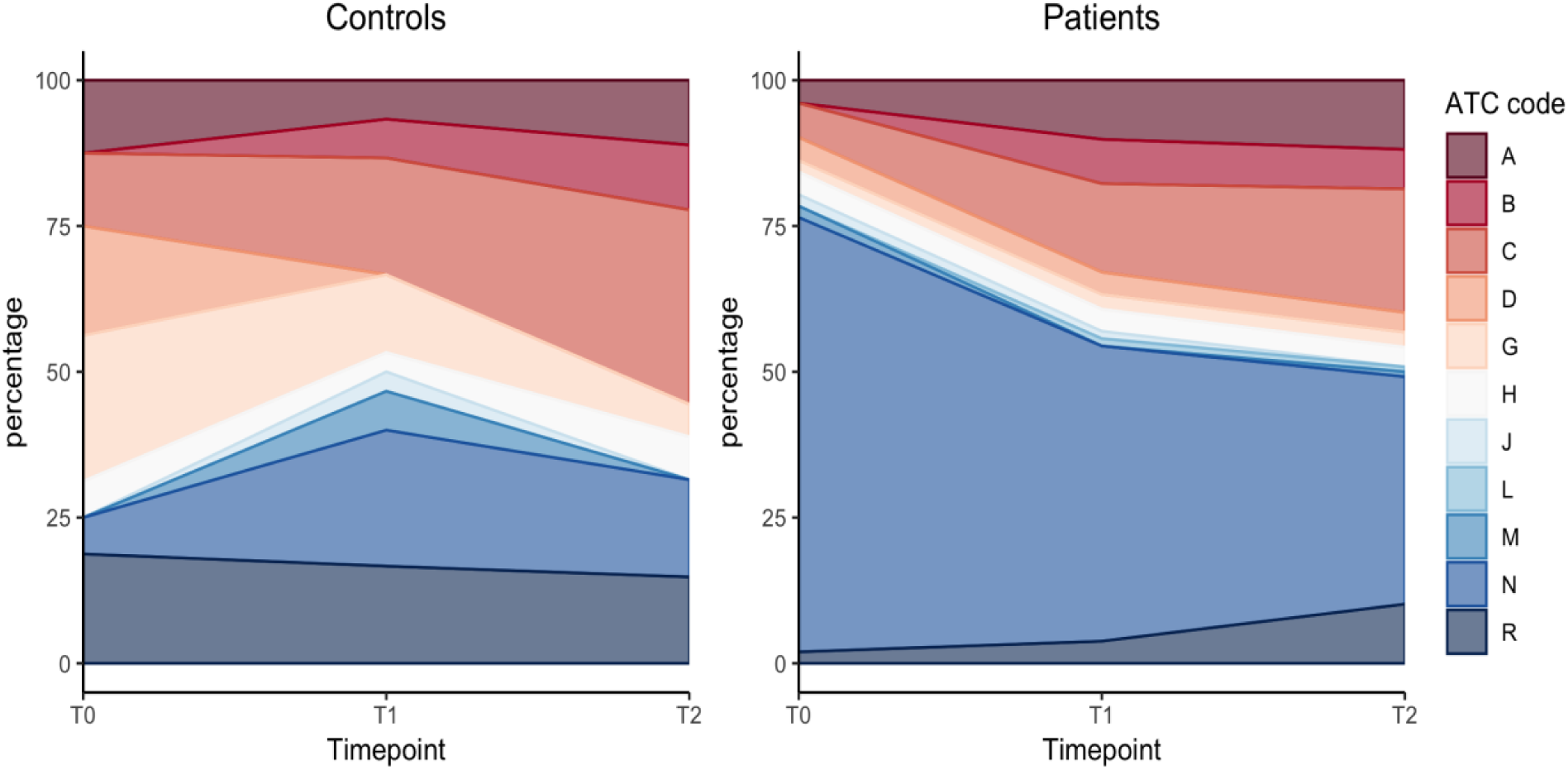
Overall use of medication in controls and patients. Area chart of percentages for each anatomical main group used at each time point based on the ATC first level. Abbreviations: ATC, anatomical therapeutic chemical classification system; A, alimentary tract and metabolism: B, blood and blood forming organs; C, cardiovascular system; D, dermatologicals; G, genito-urinary system and hormones; H, systemic hormonal preparations; J, antiinfectives for systemic use; L, antineoplastic and immunomodulating agents; M, musculo-skeletal system; N, nervous system; R, respiratory system.

### b. Individual brain tissue volume using SIENAX

Diagnostic plots of the fitted linear mixed effect model revealed one influential outlier (see supplementary material Figure S4), which was removed from further analysis. The corrected model showed a significant effect of group status on TBV (see supplementary material Table S3), with lower values in patients compared to controls by 36841.28 mm^3^ ± 11316.9 (standard errors). While the effect of time^2^ was significant, indicating a non-linear decrease in TBV over time, the time-by-group interaction was not. Hence, patients and controls showed similar trajectories in TBV atrophy over time (see Figure 3).

**Figure 3.**
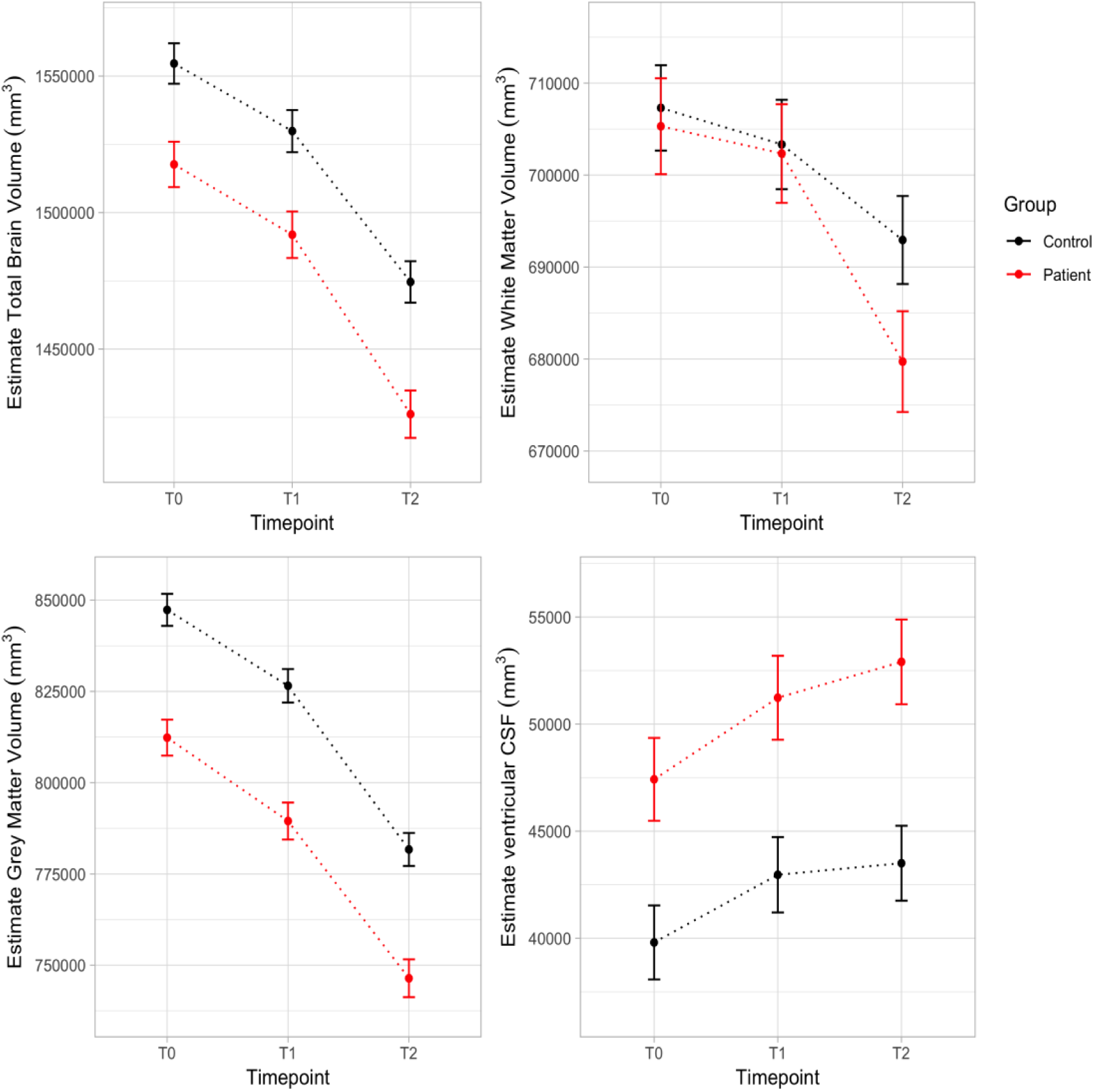
Estimated marginal means of total brain volume and tissue-specific volumes over time. Displayed are estimates for total brain volume (upper panel, left), gray matter volume (lower panel, left), ventricular cerebral spinal fluid (CSF, lower panel, left), and white matter volume (upper panel, right). For display purposes, values are based on a more parsimonious linear mixed effect model using time as a categorial factor. The model is adjusted for age at baseline, sex and body mass index. Estimates are displayed with upper and lower confidence intervals.

In line with the TBV findings, gray matter volume and ventricular CSF differed significantly between groups and over time (see Figure 3). Gray matter volume was lower and ventricular CSF was higher in patients compared to controls by 35034.29 ± 6667.44 mm^3^ (standard error) and 8557.06 ± 2620.23 mm^3^ (standard error), respectively. For both gray matter volume and CSF, we found no interaction between group and time (see Table S5 supplementary material). Contrary to the TBV results, we found a significant group x time interaction for white matter volume (Estimate=-779.52, S.E.=353.71, t=-2.20, p=0.03). Yet, white matter volume did not differ significantly between patients and controls (Estimate_Patient_=-1839.66, S.E.=7129.42, t=-0.26, p=0.80). Figure 3 (upper panel, right) shows that white matter volume is similar between groups at T0 and T1 but diverges at T2 with a higher reduction in patients relative to controls.

### c. Two-time point percentage brain and ventricular volume change using SIENA

Diagnostic plots of multiple linear regression model revealed two influential outliers for the T0-T1 and one for the T0-T2 PBVC comparisons (see supplementary material Figure S5 and S6), which were removed from analyses.

Group status, age^2^, BMI change and sex were significantly associated with PBVC from T0 to T2 (see supplementary material Table S6, Figure S7). PBVC from T0 to T2 was higher in patients relative to controls by 0.71% ± 0.25 (standard error), indicating slightly higher atrophy levels in patients over 13 years. A significant effect of group was found for T1 to T2 (Estimate_Patient_=-0.74, S.E.=0.26, t=-2.79, p=0.01), but not for PBVC from T0 to T1 (Estimate_Patient_=-0.29, S.E.=0.18, t=-1.63, p=0.11). From T0 to T2, weight gain was associated with higher brain volume loss over 13 years, across groups, and women showed a slightly higher decrease in PBVC than men, PVVC did not differ between groups for any of the two-time point comparisons (see supplementary material Table S6).

From T0 to T2, voxel-wise statistical analysis of case-control differences in brain edge points revealed significantly atrophic brain edges in the left hemisphere (see supplementary material, Figure S8), covering the parietal and temporal lobe, of patients relative to controls (contrast controls > patient, p = 0.04, peak MNI voxel: -68 -32 20). For the T0-T1 or T1-T2 comparisons, we did not find any case-control differences in atrophic brain edges.

### d. Relationships with clinical variables and SIENAX output

Repeated measures correlations (Bonferroni correction, α=0.003) showed: (1) a significant negative association between CPZ and TBV (r=-0.40, p=2.19e-04, 95% confidence interval (CI) [-0.57, -0.19]), likely driven by its association with gray matter volume (r=-0.42, p=1.08e-04, 95% CI [-0.58, -0.21]); and (2) a significant positive association between GAF scores and TBV (r=0.33, p=2.58e-03, 95% CI [0.12, 0.51]), again, likely driven by an association with gray matter volume (r=0.36, p=9.32e-04, 95% CI [0.15, 0.54]). There were no significant associations between SAPS and SANS score, and TBV after correction for multiple comparison.

We did not find any significant association between ATC-C drugs, antipsychotic class and TBV. Yet, there was a significant main effect of ATC-N drugs on TBV (Estimate=8171.84, S.E.=3223.77, t=2.54, p=0.01) and SGA on TBV (Estimate=-31568.93, S.E.=13184.63, t=-2.39, p=0.02; see supplementary material Table S7, Figure S9). While accounting for other nervous-system drug use, patients on SGA alone showed lowest TBV. Patients on a combination of FGA and SGA in addition to other nervous-system drugs seem to have higher TBV than patients on FGA or SGA alone. These results could be replicated in the model with white matter volume, but not with gray matter volume or ventricular CSF. However, when excluding ATC-N variable from the model, the main effect of SGA became marginally significant.

Controls using nervous system drugs showed a higher TBV relative to their untreated counterparts (Estimate=14719.01, S.E.=6380.10, t=2.31, p = 0.02). Here, the effect seems driven by changes in gray matter volume (Estimate=8456.24, S.E.=3612.05, t=2.34, p=0.02). We note, however, that only a limited number of controls (n = 7) used ATC-N drugs.

### a. Relationships between clinical variables and PBVC

There was no association between antipsychotic medication dosage and brain volume change, as neither CPZ at baseline (t=-1.33, p=0.19) nor cumulative CPZ (t=-0.61, p=0.55) nor CPZ change (t=0.23, p=0.82) were associated with PBVC from T0 to T2.

## Discussion

Our results suggest that patients with chronic schizophrenia show similar trajectories of aging-related brain volume loss over 13-years as healthy controls. Antipsychotic exposure did not show an association with brain volume loss longitudinally, yet, antipsychotic medication load and class, in combination with add-on drugs, may explain stable brain volume differences between patients with chronic schizophrenia.

Vita and colleagues (2012) speculated that tissue loss may be accelerated early in the illness, but less evident as the disease stabilizes^8^. In line with this claim, using SIENAX, we found baseline differences in TBV and gray matter volume as well as ventricle size in patients and controls, but no group differences in rates of volume loss over time, indicative of established and stable deficits early in the illness course. White matter volume was similar between groups during the first follow-up, but started to diverge in the second follow-up period, with higher white matter loss in patients. Using SIENA, we found greater brain volume loss in patients over 13 years; yet, again seemingly driven by emerging group differences in the second follow-up period. After excluding the first follow-up time point from the SIENAX model, the group x time interaction was significant for TBV and white matter volume. Hence, the discrepancy between SIENAX and SIENA results seems to be due to non-linear white matter volume loss in later stages of the illness. However, this result may be interpreted with cautions as white matter volume showed substantial variability between participants compared to TBV and gray matter volume.

Cross-sectional studies indicate that white matter volume differences are less pronounced, with smaller effect sizes, than gray matter differences in schizophrenia ^28^. In a twin study, genetic factors explained 77% of an association between smaller brain volume and risk for developing schizophrenia, wherein smaller white matter volume accounted for 94% of phenotypic correlations ^29^. A significant environmental correlation was found in gray matter ^29^. Our results argue against a progressive reduction in gray matter volume in chronic schizophrenia, beyond aging. However, we found a negative correlation between antipsychotic medication load, general functioning and gray matter volume. Hence, environmental influences on gray matter volume may establish early, likely based on illness severity, and remain stable over time.

Again, contrary to previous work ^14,30^, we did not find excessive ventricular enlargement in patients relative to controls over time, using both SIENA and SIENAX. Whether ventricular enlargement is a static ^29,31^ or progressive ^11,32^ neuropathological process remains debated. Here, patients had significantly larger ventricles at baseline, but the trajectories did not differ between groups. Ventricular volume appeared to show greater variation between participants, which may be consistent with previous studies suggesting both ventricular enlargement and reduction in schizophrenia ^33,34^.

We found associations between use of antipsychotic medication and brain volume. Although higher CPZ level and SGA only use was associated with lower TBV, these variables did not show an association with brain volume loss over time. These results contrast with previous work stating that antipsychotic medication is associated with brain volume loss in a dose-dependent manner ^7,35^. Yet, most studies showing dose-dependent effects of antipsychotic medication on brain morphology were conducted in first-episode patients with limited follow-up periods ^28,36^. Longitudinal studies in chronic schizophrenia are sparse. One might speculate that structural adaptations due to antipsychotic medication occur early, during the first period of treatment, and persist with no additional progressive changes over time. An exception may be clozapine. Here, patients on SGA only showed lowest TBV, with a subset of these patients being on clozapine monotherapy. Ahmed and colleagues found that patients who were switched to clozapine demonstrated reductions in brain volume and progressive cortical thinning 6–9 months after commencing treatment^38^. In addition, the add-on drugs, given to relieve side effects, may also mediate the differential impact of antipsychotic class. The effect of SGA on TBV was only evident when we accounted for other nervous-system drug use. The intake of non-antipsychotic nervous system drugs was associated with higher TBV, independent of group status. While patients were primarily on psycholeptics (e.g. lithium), psychoanaleptics (e.g. antidepressants) and anti-parkinson drugs, a limited number of controls mainly received psychoanaleptics and analgesics. Higher TBV in non-antipsychotic nervous system drug users might be particularly driven by antidepressants and lithium use, as these drugs have been linked to volumetric enlargements ^39,40^.

Another factor that influenced brain volume loss in the current study was BMI change over time. Weight gain was associated with greater brain volume loss, independent of group status, which is in line with findings in overweight ^41^ and obese populations ^42^. Still, the impact of antipsychotic class persisted after correcting for BMI, arguing for an effect of BMI independent of medication ^43^. However, we found higher weight gain in female relative to male patients over 13 years. Female sex and polypharmacy are seemingly risk factors for reporting a severe side effect burden ^44^. Women with psychotic disorders show an increased susceptibility to weight gain, diabetes, and specific cardiovascular risks of antipsychotics ^45^. Yet, the study of long-term consequences of antipsychotic medication for women’s brain health has been neglected in basic and preclinical research. Here, we found that female patients had larger brain volume change over 13 years than male patients, most likely driven by their higher BMI. The further assessment of sex-specific medication effects was not possible due to the small sample of women in the study. Future studies should ensure sex-specific reporting of research findings to improve clinical outcomes for both men and women.

Whilst it is of importance to assess causes of brain volume loss in schizophrenia, of equal importance is the clinical significance of change in brain morphology. Here, we found that global functioning was positively correlated with TBV, indicating that larger TBV constitutes better general functioning. General functioning evaluated with GAF also showed a negative association with CPZ (r=-0.254, p=0.03, 95% CI [-0.457, -0.025]), indicating that low-functioning patients are prescribed higher doses of antipsychotics. Negative symptoms in patients also increased with age but we did not find any significant effect between SANS scores and TBV after correction for multiple comparison.

The major strength of the current study is the long follow-up period and the careful and detailed assessment of medication use at each time point for patients and controls. In addition, alcohol dependencies and illicit drug use did not differ significantly between patients and healthy controls at baseline, and are therefore unlikely to influence the current findings. However, we also acknowledge a number of limitations. A major limitation is the modest sample size with a largely unbalanced sex distribution, which preluded the robust analysis of sex differences, diagnostic subgroup differences and the impact of individual drugs on brain volume change. Another limitation is the change in scanner from T1 to T2. Scanner hardware and software changes are a critical issue in longitudinal neuroimaging studies. To circumvent the confound of scanner change, we used SIENA, a superior image analysis method that has proven to be robust to changes in acquisition protocols ^46^, and entered scanner as fixed factor to every statistical model using SIENAX data. Although extensive, medication data was restricted to the three time points. Thus, we lack the temporal resolution to determine when patients might have changed between antipsychotic medication types (e.g. from FGA to SGA, or switch to polypharmacy) and how long they were taking certain medications. Further, the distinction between the categories FGA and SGA has, although widely used, questionable validity ^47^.

In summary, patients with chronic and long-term treated schizophrenia showed lower total and tissue-specific brain volume at baseline compared to healthy controls. However, there was limited evidence of progressive brain volume loss over 13-years. Antipsychotic medication and other nervous-system drugs such as antidepressants and mood stabilizers may contribute differentially to brain volume changes in patients, highlighting the need to assess add-on medication.

## Acknowledgements

We thank the study participants and the clinicians involved in the recruitment and assessment. A special thanks to Monica Hellberg, Charlotta Leandersson and Sara Holmqvist for technical assistance. This work was supported by the Swedish Research Council (grant numbers: 2003-5845, 2006-2992, 2006-986, 2008-2167, K2012-61X-15078-09-3, 521-2011-4622, 521-2014-3487, 2017-00949) and the South-Eastern Norway Regional Health Authority (grant number: 2017093, 2017-097).

## Author Contributions

CB undertook the processing of the imaging data, the statistical analysis, the literature search, interpreted the results, wrote the first draft and finalized the manuscript. KNJ and SN helped with the processing of the imaging data. KNJ also contributed during the statistical analysis and interpretation of the results. IA and EGJ designed the longitudinal HUBIN study and obtained funding. EGJ performed the clinical interviews. KNJ, LW, SN, IA and EGJ critically revised the first draft and approved the final manuscript.

## References

1. James SL, Abate D, Abate KH, et al. Global, regional, and national incidence, prevalence, and years lived with disability for 354 diseases and injuries for 195 countries and territories, 1990-2017: a systematic analysis for the Global Burden of Disease Study 2017. Lancet. 2018;392(10159):1789–1858.

2. Hulshoff Pol HE, Kahn RS. What happens after the first episode? A review of progressive brain changes in chronically ill patients with schizophrenia. Schizophr Bull. 2008;34(2):354–366.

3. Olabi B, Ellison-Wright I, McIntosh AM, Wood SJ, Bullmore E, Lawrie SM. Are there progressive brain changes in schizophrenia? A meta-analysis of structural magnetic resonance imaging studies. Biol Psychiatry. 2011;70(1):88–96.

4. Andreasen NC, Nopoulos P, Magnotta V, Pierson R, Ziebell S, Ho BC. Progressive brain change in schizophrenia: a prospective longitudinal study of first-episode schizophrenia. Biol Psychiatry. 2011;70(7):672–679.

5. Cannon TD, Chung Y, He G, et al. Progressive reduction in cortical thickness as psychosis develops: a multisite longitudinal neuroimaging study of youth at elevated clinical risk. Biol Psychiatry. 2015;77(2):147–157.

6. Gutierrez-Galve L, Chu EM, Leeson VC, et al. A longitudinal study of cortical changes and their cognitive correlates in patients followed up after first-episode psychosis. Psychol Med. 2015;45(1):205–216.

7. Veijola J, Guo JY, Moilanen JS, et al. Longitudinal changes in total brain volume in schizophrenia: relation to symptom severity, cognition and antipsychotic medication. PLoS One. 2014;9(7):e101689.

8. Vita A, De Peri L, Deste G, Sacchetti E. Progressive loss of cortical gray matter in schizophrenia: a meta-analysis and meta-regression of longitudinal MRI studies. Transl Psychiatry. 2012;2:e190.

9. Schaufelberger MS, Lappin JM, Duran FL, et al. Lack of progression of brain abnormalities in first-episode psychosis: a longitudinal magnetic resonance imaging study. Psychol Med. 2011;41(8):1677–1689.

10. Haukvik UK, Hartberg CB, Nerland S, et al. No progressive brain changes during a 1-year follow-up of patients with first-episode psychosis. Psychol Med. 2016;46(3):589–598.

11. Fusar-Poli P, Smieskova R, Kempton MJ, Ho BC, Andreasen NC, Borgwardt S. Progressive brain changes in schizophrenia related to antipsychotic treatment? A meta-analysis of longitudinal MRI studies. Neurosci Biobehav Rev. 2013;37(8):1680–1691.

12. Ho BC, Andreasen NC, Ziebell S, Pierson R, Magnotta V. Long-term antipsychotic treatment and brain volumes: a longitudinal study of first-episode schizophrenia. Arch Gen Psychiatry. 2011;68(2):128–137.

13. Lieberman JA, Tollefson GD, Charles C, et al. Antipsychotic drug effects on brain morphology in first-episode psychosis. Arch Gen Psychiatry. 2005;62(4):361–370.

14. Huhtaniska S, Jaaskelainen E, Hirvonen N, et al. Long-term antipsychotic use and brain changes in schizophrenia - a systematic review and meta-analysis. Hum Psychopharmacol. 2017;32(2).

15. Vares M, Saetre P, Stralin P, Levander S, Lindstrom E, Jonsson EG. Concomitant medication of psychoses in a lifetime perspective. Hum Psychopharmacol. 2011;26(4-5):322–331.

16. Andreasen NC. Scale for the assessment of negative symptoms (SANS). University of Iowa, Iowa City. 1983.

17. Andreasen NC. The Scale for the assessment of Positive Symptoms (SAPS). University of Iowa, Iowa City. 1984.

18. Association. AP. Diagnostic and Statistical Manual of Mental Disorders, Third Edition - Revised. Washington DC: American Psychiatric Association. 1987.

19. Spitzer RL, Williams JBW, Gibbon M, First MB. Structured Clinical Interview for DSM-III-R - Patient Version (SCID-P).. New York: Biometrics Research Department, New York State Psychiatric Institute. 1988.

20. Andreasen NC, Pressler M, Nopoulos P, Miller D, Ho BC. Antipsychotic dose equivalents and dose-years: a standardized method for comparing exposure to different drugs. Biol Psychiatry. 2010;67(3):255–262.

21. Methodology WCCfDS. Guidelines for ATC classification and DDD assignment 2019. Oslo, Norway. 2018.

22. Smith SM, De Stefano N, Jenkinson M, Matthews PM. Normalized accurate measurement of longitudinal brain change. J Comput Assist Tomo. 2001;25(3):466–475.

23. Smith SM, Zhang YY, Jenkinson M, et al. Accurate, robust, and automated longitudinal and cross-sectional brain change analysis. Neuroimage. 2002;17(1):479–489.

24. Bates D, Machler M, Bolker BM, Walker SC. Fitting Linear Mixed-Effects Models Using lme4. J Stat Softw. 2015;67(1):1–48.

25. Kuznetsova A, Brockhoff PB, Christensen RHB. lmerTest Package: Tests in Linear Mixed Effects Models. J Stat Softw. 2017;82(13):1–26.

26. Chung YJ, Rabe-Hesketh S, Dorie V, Gelman A, Liu JC. A Nondegenerate Penalized Likelihood Estimator for Variance Parameters in Multilevel Models. Psychometrika. 2013;78(4):685–709.

27. Bakdash JZ, Marusich LR. Repeated Measures Correlation. Front Psychol. 2017;8.

28. Haijma SV, Van Haren N, Cahn W, Koolschijn PC, Hulshoff Pol HE, Kahn RS. Brain volumes in schizophrenia: a meta-analysis in over 18 000 subjects. Schizophr Bull. 2013;39(5):1129–1138.

29. van Haren NE, Rijsdijk F, Schnack HG, et al. The genetic and environmental determinants of the association between brain abnormalities and schizophrenia: the schizophrenia twins and relatives consortium. Biol Psychiatry. 2012;71(10):915–921.

30. Guo JY, Huhtaniska S, Miettunen J, et al. Longitudinal regional brain volume loss in schizophrenia: Relationship to antipsychotic medication and change in social function. Schizophr Res. 2015;168(1-2):297–304.

31. de Zwarte SMC, Brouwer RM, Agartz I, et al. The Association Between Familial Risk and Brain Abnormalities Is Disease Specific: An ENIGMA-Relatives Study of Schizophrenia and Bipolar Disorder. Biol Psychiatry. 2019;86(7):545–556.

32. Kempton MJ, Stahl D, Williams SC, DeLisi LE. Progressive lateral ventricular enlargement in schizophrenia: a meta-analysis of longitudinal MRI studies. Schizophr Res. 2010;120(1-3):54–62.

33. DeLisi LE, Hoff AL, Kushner M, Calev A, Stritzke P. Left ventricular enlargement associated with diagnostic outcome of schizophreniform disorder. Biol Psychiatry. 1992;32(2):199–201.

34. Puri BK, Hutton SB, Saeed N, et al. A serial longitudinal quantitative MRI study of cerebral changes in first-episode schizophrenia using image segmentation and subvoxel registration. Psychiat Res-Neuroim. 2001;106(2):141–150.

35. Andreasen NC, Liu D, Ziebell S, Vora A, Ho BC. Relapse duration, treatment intensity, and brain tissue loss in schizophrenia: a prospective longitudinal MRI study. Am J Psychiatry. 2013;170(6):609–615.

36. Emsley R, Asmal L, du Plessis S, Chiliza B, Phahladira L, Kilian S. Brain volume changes over the first year of treatment in schizophrenia: relationships to antipsychotic treatment. Psychol Med. 2017;47(12):2187–2196.

37. van Haren NE, Schnack HG, Cahn W, et al. Changes in cortical thickness during the course of illness in schizophrenia. Arch Gen Psychiatry. 2011;68(9):871–880.

38. Ahmed M, Cannon DM, Scanlon C, et al. Progressive Brain Atrophy and Cortical Thinning in Schizophrenia after Commencing Clozapine Treatment. Neuropsychopharmacology. 2015;40(10):2409–2417.

39. Dusi N, Barlati S, Vita A, Brambilla P. Brain Structural Effects of Antidepressant Treatment in Major Depression. Curr Neuropharmacol. 2015;13(4):458–465.

40. Lyoo IK, Dager SR, Kim JE, et al. Lithium-induced gray matter volume increase as a neural correlate of treatment response in bipolar disorder: a longitudinal brain imaging study. Neuropsychopharmacology. 2010;35(8):1743–1750.

41. Gupta A, Mayer EA, Sanmiguel CP, et al. Patterns of brain structural connectivity differentiate normal weight from overweight subjects. Neuroimage-Clin. 2015;7:506–517.

42. Raji CA, Ho AJ, Parikshak NN, et al. Brain structure and obesity. Hum Brain Mapp. 2010;31(3):353–364.

43. Bobb JF, Schwartz BS, Davatzikos C, Caffo B. Cross-sectional and longitudinal association of body mass index and brain volume. Hum Brain Mapp. 2014;35(1):75–88.

44. Iversen TSJ, Steen NE, Dieset I, et al. Side effect burden of antipsychotic drugs in real life - Impact of gender and polypharmacy. Prog Neuropsychopharmacol Biol Psychiatry. 2018;82:263–271.

45. Seeman MV. Secondary effects of antipsychotics: women at greater risk than men. Schizophr Bull. 2009;35(5):937–948.

46. de Bresser J, Portegies MP, Leemans A, Biessels GJ, Kappelle LJ, Viergever MA. A comparison of MR based segmentation methods for measuring brain atrophy progression. Neuroimage. 2011;54(2):760–768.

47. Leucht S, Corves C, Arbter D, Engel RR, Li C, Davis JM. Second-generation versus first-generation antipsychotic drugs for schizophrenia: a meta-analysis. Lancet. 2009;373(9657):31–41.

48. Allen M, Poggiali D, Whitaker K, Marshall TR, Kievit RA. Raincloud plots: a multi-platform tool for robust data visualization. Wellcome Open Res. 2019;4:63.

