## Supplementary Material for "Trajectories of brain volume change over 13 years in chronic schizophrenia"

### Methods

#### *Participants*

For study inclusion at baseline, participants were required to have no history of head trauma resulting in loss of consciousness for more than 5 minutes and no history of somatic illness which could affect brain structure. Somatic illness was excluded by a thorough physical investigation and biochemical screening. Control participants with severe mental disorders among first-degree relatives were excluded. Lifetime psychiatric diagnoses were established according to Diagnostic and Statistical Manual of Mental Disorder (DSM)-IIIR or DSM-IV based on hospital case notes and structured clinical interviews performed by trained staff [1]. Detailed inclusion procedures are described elsewhere [2].

#### *MRI data acquisition*

MRI data at baseline and ~5-year follow-up was collected using a General Electric 1.5-Tesla Signa HDxt scanner equipped with an 8-channel head coil. T1-weighted volumes were acquired using a three-dimensional spoiled gradient recalled (SPGR) pulse sequence with the following parameters: 124 coronal slices, flip angle = 35°, repetition time = 24 ms, echo time = 6.0 ms, acquisition matrix = 256 x 192, field of view = 24 cm. At ~13-year follow-up, MRI images were acquired at General Electric 3-Tesla Discovery MR750 scanner equipped with a 32-channel head coil. Imaging data was acquired using a three-dimensional gradient echo sequence with the following parameters: sagittal orientation, flip angle = 12°, repetition time = 7.90 ms, echo time = 3.1 ms, inversion time = 450 ms, slice thickness = 1.2 mm, bandwidth = 244.14 Hz/pixel, voxel size = 1 x 1 x 1.2 mm.

#### *SIENAX – Cross-sectional pipeline*

SIENAX was used to calculate individual total brain volume (TBV) and tissue-specific volumes of grey matter, white matter and ventricular cerebral spinal fluid (CSF) at each time point. Measures were normalized for participant head size [3,4]. SIENAX starts by extracting brain and skull images from the single whole-head input data [5]. To best optimize brain extraction for our dataset, we pre-masked the abovementioned images. In detail, we used (i) *fslorient2std* to reorient the whole-head input data to match the orientation of the standard template images, (ii) *standard_space_roi -b* was applied to generate a more generous and robust mask than the default brain extraction command, and (iii) we determined the voxel coordinates of the center of gravity on the generated output using *fslstats -C* and fed them to SIENAX brain extraction via the *-c* parameter. The optimized brain extraction pipeline was subsequently applied to the original image. Further, we found a fractional intensity threshold of 0.35 to be optimal for our dataset. The brain image is then affine-registered to MNI152 space [6,7] (using the skull image to determine the registration scaling); this is primarily in order to obtain the volumetric scaling factor, to be used as a normalization for head size. Next, tissue-type segmentation with partial volume estimation is carried out [8] in order to calculate total volume of brain tissue.

***SIENA – Longitudinal pipeline***

SIENA was used to estimate two-time point percentage brain volume change (PBVC) and percentage ventricular volume changes (PVVC) from T0 to T1, T1 to T2 and T0 to T2 [3,4], to cross-valid SIENAX results. Before running SIENA, the image dimensions of MRI images at T0 and T1, acquired at 1.5T (256 x 256 x 124 $mm^3$), were conformed to T2 images, acquired at 3T (146 x 256 x 256 $mm^3$), by adding 22 slices in z-dimension (256 x 256 x 146 $mm^3$) using *fslroi* (FSL) to normalize differences and to optimize registration.

SIENA starts by extracting brain and skull images from the two-time point whole-head input data [5]. We applied the same optimized brain extraction pipeline mentioned above to SIENA. The two extracted brain images were then aligned to each other [6,7] (using the skull images to constrain the registration scaling); both brain images were resampled into the space halfway between the two. Next, tissue-type segmentation was carried out [8] in order to find brain/non-brain edge points, and then perpendicular edge displacement (between the two time points) was estimated at these edge points. Finally, the mean edge displacement was converted into a (global) estimate of percentage brain volume change between the two time points. SIENA outputs were visually inspected, without knowledge of clinical and demographic characteristics, to ensure data quality. To calculate PVVC, extension for ventricular analysis using the *-V* parameter was activated.

For computing voxel wise multi-participant statistics using the generated SIENA output, the edge displacement image (encoding, at brain/non-brain edge points, the outwards or inwards edge change between the two time points) was dilated, transformed into MNI152 space, and masked by a standard MNI152-space brain edge image on a single participant level. In this way the edge displacement values were warped onto the standard brain edge [9]. Next, the resulting images from all participants were fed into voxel wise statistical analysis to identify brain edge points which are significantly atrophic between groups.

***Statistical analysis***

***a. Demographic and clinical data***

To examine the number of 1st level anatomical therapeutic chemical classification system (ATC) co-medication per participant as a function of group and time, we fitted Bayesian generalized linear mixed-effects models with Poisson distribution using blme (details main text). For all models except of one no priors were imposed over the fixed effect. However, modelling 1st level ATC D code (Dermatologicals), we set a normal-prior on the fixed effects to handle complete separation due to zero observations in the control group at T0 and T1. We were not able to fit a model for 1st level ATC L code (Antineoplastic and immunomodulating agents) due to limited data points (see Table 3, main text).

### Results

#### *Tables*

#### Table S1| Demographics and clinical characteristics of included and excluded participants.

| | Control | | | | Patient | | | |
|---|---|---|---|---|---|---|---|---|
| | **Excluded** | **Included** | **p** | **test** | **Excluded** | **Included** | **p** | **test** |
| **N** | 47 | 79 | | | 63 | 64 | | |
| **Sex, women N (%)** | 16 (34.0) | 29 (36.7) | 0.913 | $\chi_2$ | 21 (33.3) | 13 (20.3) | 0.145 | $\chi_2$ |
| **Age at scan (y)** | 39.7 [17.9] | 46.2 [9.0] | **0.019** | KW | 43.3 [10.3] | 42.6 [9.9] | 0.504 | KW |
| **Handedness (%)** | | | 1.000 | FET | | | 1.000 | FET |
| *right* | 40 (93.0) | 70 (89.7) | | | 49 (84.5) | 51 (83.6) | | |
| *left* | 2 (4.7) | 5 (6.4) | | | 6 (10.3) | 7 (11.5) | | |
| *ambidextrous* | 1 (2.3) | 3 (3.8) | | | 3 (5.2) | 3 (4.9) | | |
| **Education (y)** | 14.0 [4.0] | 14.0 [4.0] | 0.938 | KW | 12.0 [4.0] | 12.0 [3.0] | 0.132 | KW |
| **SES Father** | 5.0 [4.0] | 5.0 [4.0] | 0.350 | KW | 5.0 [5.0] | 5.0 [5.0] | 0.739 | KW |
| **SES Mother** | 5.0 [3.0] | 5.0 [6.5] | 0.743 | KW | 5.0 [5.0] | 5.0 [5.0] | 0.970 | KW |
| **Illicit drug use, N (%)** | | | 0.604 | FET | | | 0.316 | FET |
| *never* | 22 (59.5) | 42 (53.2) | | | 21 (44.7) | 23 (47.9) | | |
| *up to 10 times* | 13 (35.1) | 25 (31.6) | | | 17 (36.2) | 16 (33.3) | | |
| *11 to 100 times* | 1 (2.7) | 7 (8.9) | | | 2 (4.3) | 6 (12.5) | | |
| *> 100 times* | 1 (2.7) | 5 (6.3) | | | 7 (14.9) | 3 (6.2) | | |
| **Alcohol use, N (%)** | | | 0.319 | FET | | | 0.450 | FET |
| *none* | 35 (97.2) | 72 (92.3) | | | 39 (79.6) | 42 (87.5) | | |
| *abuse* | 0 (0.0) | 5 (6.4) | | | 4 (8.2) | 1 (2.1) | | |
| *dependence* | 1 (2.8) | 1 (1.3) | | | 6 (12.2) | 5 (10.4) | | |
| **Age of Onset (y)** | | | | | 25.3 [9.6] | 23.5 [5.5] | 0.792 | KW |
| **Duration of illness (y)** | | | | | 16.9 (9.6) | 16.2 (8.1) | 0.653 | t-test |
| **SANS, composite score** | | | | | 21.0 [13.5] | 20.0 [15.5] | 0.509 | KW |
| **SAPS, composite score** | | | | | 8.0 [12.0] | 7.0 [10.7] | 0.815 | KW |
| **GAF, Axis 5** | | | | | 50.0 [10.00] | 50.0 [20.0] | 0.606 | KW |

* distributed data in mean (standard deviation), non-normal distributed data in median [Interquartile range],

Abbreviations: N, number; y, year; IQ, intelligence quotient; SES, socio-economic status; KW, Kruskal-Wallis; FET, Fisher exact test; SANS, scale for the assessment of negative symptoms; SAPS, scale for the assessment of positive symptoms; GAF, global assessment of functioning.

**Table S2| (Co)administration of other non-antipsychotic drugs over time using ATC first-level classification.**

| | | Controls | | | Patients | | |
|---|---|---|---|---|---|---|---|
| | | **T0 (N)** | **T1 (N)** | **T2 (N)** | **T0 (N)** | **T1 (N)** | **T2 (N)** |
| **ATC, 1st level*** | No medication (except antipsychotics) | 65 | 39 | 35 | 34 | 20 | 13 |
| **Code** | **Content** | | | | | | |
| **A** | Alimentary tract and metabolism | 2 | 2 | 6 | 2 | 8 | 14 |
| **B** | Blood and blood forming organs | 0 | 2 | 6 | 0 | 6 | 8 |
| **C** | Cardiovascular system | 2 | 6 | 18 | 3 | 12 | 25 |
| **D** | Dermatologicals | 3 | 0 | 0 | 2 | 3 | 4 |
| **G** | Genito-urinary system and sex hormones | 4 | 4 | 3 | 1 | 2 | 3 |
| **H** | Systemic hormonal preparations | 1 | 1 | 4 | 2 | 3 | 4 |
| **J** | Antiinfectives for systemic use | 0 | 1 | 0 | 1 | 1 | 0 |
| **L** | Antineoplastic and immunomodulating agents | 0 | 0 | 0 | 0 | 1 | 1 |
| **M** | Musculo-skeletal system | 0 | 2 | 0 | 1 | 0 | 1 |
| **N** | Nervous system | 1 | 7 | 9 | 38 | 40 | 46 |
| **R** | Respiratory system | 3 | 5 | 8 | 1 | 3 | 12 |

*****defined as medication given on a daily basis, Abbreviation: ATC, anatomical therapeutic chemical classification system; N, number; T0, baseline; T1, first follow-up; T2, second follow-up.

**Table S3| Second-level ATC classification for selected main groups.**

| | | | Control | | | Patient | | |
|---|---|---|---|---|---|---|---|---|
| ATC, 2nd level | | | T0 | T1 | T2 | T0 | T1 | T2 |
| Main group | Code | Therapeutic subgroup | N | N | N | N | N | N |
| Cardiovascular system | 01 | Cardiac therapy | 0 | 0 | 0 | 0 | 0 | 1 |
| | 03 | Diuretics | 0 | 0 | 1 | 1 | 1 | 4 |
| | 07 | Beta blocking agents | 1 | 2 | 3 | 1 | 4 | 7 |
| | 08 | Calcium channel blockers | 0 | 0 | 2 | 0 | 1 | 3 |
| | 09 | Agents acting on the renin–angiotensin system | 1 | 3 | 7 | 0 | 3 | 4 |
| | 10 | Lipid modifying agents | 0 | 1 | 5 | 1 | 3 | 6 |
| Nervous system | 02 | Analgesics | 0 | 2 | 2 | 0 | 0 | 1 |
| | 03 | Antiepileptics | 0 | 0 | 0 | 1 | 2 | 5 |
| | 04 | Anti-parkinson drugs | 0 | 0 | 1 | 11 | 9 | 8 |
| | 05 | Psycholeptics | 0 | 1 | 1 | 19 | 22 | 16 |
| | 06 | Psychoanaleptics | 1 | 4 | 5 | 7 | 7 | 14 |
| | 07 | Other nervous system drugs | 0 | 0 | 0 | 0 | 0 | 1 |

* *only agents with counts are listed;* Abbreviations: ATC, anatomical therapeutic chemical classification system; N, number; T0, baseline; T1, ~5-year follow-up; T2, ~13-year follow-up.

**Table S4| Summary of SIENAX total brain volume linear mixed effect model, with and without outlier.**

| **Fixed Effects** | **Estimate** | **S.E.** | **t-value** | **p-values** |
|---|---|---|---|---|
| **Including outlier** | | | | |
| Group (Patient) | -38272.14 | 11817.39 | - 3.24 | **1.47e-03** |
| Time (continuous) | -655.96 | 1725.39 | -0.38 | 0.70 |
| $Time_2$ | -7327.25 | 27987.99 | -2.62 | **9.42e-03** |
| Age (at baseline) | -5371.24 | 5837.28 | -0.92 | 0.36 |
| $Age_2$ | 142.24 | 755.83 | 0.19 | 0.85 |
| Sex | 15711.85 | 12186.00 | 1.29 | 0.20 |
| BMI | -1115.14 | 785.72 | -1.42 | 0.16 |
| Scanner | 54180.98 | 29163.07 | 1.86 | *0.07* |
| Group (Patient) x Time Interaction | -1059.87 | 564.41 | -1.88 | *0.06* |
| **Random Effects** | **Variance** | **Std. Dev.** | | |
| Participant (Intercept) | 3844273571.74 | 62002.21 | | |
| Residuals | 767118823.60 | 27696.91 | | |
| **Excluding outlier** | | | | |
| Group (Patient) | -36841.28 | 11316.91 | -3.26 | **1.39e-03** |
| Time (continuous) | -124.93 | 1653.24 | -0.08 | 0.94 |
| $Time_2$ | -8460.65 | 2684.05 | -3.15 | **1.84e-03** |
| Age (at baseline) | -4360.59 | 5578.34 | -0.78 | 0.44 |
| $Age_2$ | 19.23 | 722.21 | 0.03 | 0.98 |
| Sex | 14253.63 | 11639.75 | 1.23 | 0.22 |
| BMI | -877.20 | 751.97 | -1.17 | 0.24 |
| Scanner | 66724.34 | 27980.94 | 2.39 | **0.02** |
| Group (Patient) x Time Interaction | -779.36 | 542.94 | -1.44 | 0.15 |
| **Random Effects** | **Variance** | **Std. Dev.** | | |
| Participant (Intercept) | 3.502e+09 | 59177 | | |
| Residuals | 7.015e+08 | 26486 | | |

Abbreviations: S.E., standard error; Std. Dev., standard deviation; BMI, body mass index.

**Table S5| Summary of SIENAX tissue-specific linear mixed effect model.**

| Fixed Effects | Estimate | S.E. | t-value | p-values |
|---|---|---|---|---|
| **Gray Matter Volume** | | | | |
| Group (Patient) | -35034.29 | 6667.44 | -5.26 | 4.67e-07 |
| Time (continuous) | -4078.63 | 1052.95 | -3.87 | 1.40e-04 |
| $Time_2$ | -297.84 | 1708.74 | -0.17 | 0.86 |
| Age (at baseline) | -6358.06 | 3265.38 | -1.95 | *0.05* |
| $Age_2$ | 313.74 | 422.76 | 0.74 | 0.46 |
| Sex | 23066.90 | 6812.62 | 3.39 | **9.28e-04** |
| BMI | -1446.18 | 463.60 | -3.12 | **1.96e-03** |
| Scanner | -7158.90 | 17818.86 | -0.40 | 0.69 |
| Group (Patient) x Time Interaction | -3.33 | 346.67 | -0.01 | 0.99 |
| **Random Effects** | **Variance** | **Std. Dev.** | | |
| Participant (Intercept) | 1180075646.40 | 34352.23 | | |
| Residuals | 286434194.70 | 16924.37 | | |
| **White Matter Volume** | | | | |
| Group (Patient) | -1839.66 | 7129.42 | -0.26 | 0.80 |
| Time (continuous) | 3928.67 | 1075.97 | 3.65 | **3.24e-04** |
| $Time_2$ | -8124.49 | 1746.56 | -4.65 | **5.58e-06** |
| Age (at baseline) | 1978.33 | 3505.25 | 0.56 | 0.57 |
| $Age_2$ | -291.90 | 453.82 | -0.64 | 0.52 |
| Sex | -8787.46 | 7313.67 | -1.20 | 0.23 |
| BMI | 621.02 | 483.02 | 1.29 | 0.20 |
| Scanner | 73578.09 | 18209.87 | 4.04 | **7.30e-05** |
| Group (Patient) x Time Interaction | -779.52 | 353.71 | -2.20 | **0.03** |
| **Random Effects** | **Variance** | **Std. Dev.** | | |
| Participant (Intercept) | 1373693717.3 | 37063.37 | | |
| Residuals | 297916853.6 | 17260.27 | | |
| **Ventricular volume** | | | | |
| Group (Patient) | 8557.06 | 2620.23 | 3.27 | **1.36e-03** |
| Time (continuous) | 37.44 | 268.20 | 0.14 | 0.89 |
| $Time_2$ | 1102.72 | 435.87 | 2.53 | **0.01** |

| | | | | |
|---|---|---|---|---|
| Age (at baseline) | -1851.94 | 1316.26 | -1.41 | 0.16 |
| Age2 | 327.49 | 170.41 | 1.92 | *0.06* |
| Sex | -3960.76 | 2747.22 | -1.44 | 0.15 |
| BMI | 217.24 | 135.70 | 1.60 | 0.11 |
| Scanner | -15771.36 | 4539.86 | -3.47 | **6.18e-04** |
| Group (Patient) x Time Interaction | 136.09 | 87.44 | 1.56 | 0.12 |
| **Random Effects** | **Variance** | **Std. Dev.** | | |
| Participant (Intercept) | 204043136.58 | 14284.37 | | |
| Residuals | 18116780.19 | 4256.38 | | |

Ab Abbreviations: S.E., standard error; Std. Dev., standard deviation; BMI, body mass index.

### Table S6| Multiple linear regression model for two-timepoint percent brain and ventricular volume change comparisons.

| Model | Estimate | S.E. | t value | p value | F | df | p | *Adj. $R^2$* |
|---|---|---|---|---|---|---|---|---|
| **PBVC** | | | | | 5.18 | 97 | **4.92e-05** | **0.22** |
| **T0-T1** | | | | | | | | |
| Group (Patient) | -0.29 | 0.18 | -1.63 | 0.11 | | | | |
| BMI change | -0.10 | 0.04 | -2.53 | **0.01** | | | | |
| Age at T0 | 0.16 | 0.10 | 1.55 | 0.13 | | | | |
| Age2 | -0.03 | 0.01 | -2.03 | *0.05* | | | | |
| ISI | 17.10 | 9.77 | 1.75 | *0.08* | | | | |
| ISI2 | -16.26 | 9.13 | -1.78 | *0.08* | | | | |
| Sex | -0.09 | 0.19 | -0.49 | 0.63 | | | | |
| **T1-T2** | | | | | 4.49 | 70 | **3.53e-04** | **0.24** |
| Group (Patient) | -0.74 | 0.26 | -2.79 | **0.01** | | | | |
| BMI change | -0.08 | 0.05 | -1.47 | 0.15 | | | | |
| Age at T0 | 0.08 | 0.16 | 0.50 | 0.62 | | | | |
| Age2 | -0.02 | 0.02 | -0.82 | 0.41 | | | | |
| ISI | -8.07 | 3.20 | -2.53 | **0.01** | | | | |
| ISI2 | 4.74 | 1.99 | 2.38 | **0.02** | | | | |
| Sex | -0.45 | 0.29 | -1.54 | 0.13 | | | | |
| **T0-T2** | | | | | 8.88 | 97 | **2.08e-08** | **0.35** |

| | | | | | | | | |
|---|---|---|---|---|---|---|---|---|
| Group (Patient) | -0.71 | 0.25 | -2.79 | | | | | |
| BMI change | -0.16 | 0.04 | -4.05 | | | | | |
| Age at T0 | 0.18 | 0.12 | 1.46 | | | | | |
| Age$_{2}$ | -0.03 | 0.02 | -2.12 | | | | | |
| ISI | -9.99 | 6.38 | -1.57 | | | | | |
| ISI$_{2}$ | 3.54 | 2.42 | 1.46 | | | | | |
| Sex | -0.52 | 0.25 | -2.03 | | | | | |
| **PVVC** | | | | | | | | |
| **T0-T1** | | | | | 5.57 | 97 | **2.04e-05** | **0.24** |
| Group (Patient) | 1.78 | 1.58 | 1.13 | 0.26 | | | | |
| BMI change | 0.08 | 0.35 | 0.24 | 0.81 | | | | |
| Age at T0 | -1.52 | 0.92 | -1.66 | 0.10 | | | | |
| Age$_{2}$ | 0.26 | 0.12 | 2.19 | **0.03** | | | | |
| ISI | -153.49 | 87.59 | -1.75 | *0.08* | | | | |
| ISI$_{2}$ | 146.85 | 81.85 | 1.79 | *0.08* | | | | |
| Sex | -3.53 | 1.74 | -2.03 | **0.05** | | | | |
| **T1-T2** | | | | | 4.04 | 69 | **9.17e-04** | **0.22** |
| Group (Patient) | 0.78 | 2.88 | 0.27 | 0.79 | | | | |
| BMI change | -0.23 | 0.57 | -0.39 | 0.70 | | | | |
| Age at T0 | 0.18 | 1.77 | 0.10 | 0.92 | | | | |
| Age$_{2}$ | 0.09 | 0.23 | 0.40 | 0.69 | | | | |
| ISI | -13.44 | 34.52 | -0.39 | 0.70 | | | | |
| ISI$_{2}$ | 9.67 | 21.53 | 0.45 | 0.66 | | | | |
| Sex | -5.52 | 3.19 | -1.73 | *0.09* | | | | |
| **T0-T2** | | | | | 9.15 | 97 | **1.22e-08** | **0.35** |
| Group (Patient) | -0.05 | 3.04 | -0.02 | 0.99 | | | | |
| BMI change | 0.57 | 0.46 | 1.22 | 0.23 | | | | |
| Age at T0 | -0.01 | 1.49 | -3.94e-03 | 1.00 | | | | |
| Age$_{2}$ | 0.15 | 0.19 | 0.77 | 0.44 | | | | |
| ISI | -39.77 | 76.45 | -0.52 | 0.60 | | | | |
| ISI$_{2}$ | 16.83 | 29.04 | 0.58 | 0.56 | | | | |
| Sex | -8.69 | 3.05 | -2.85 | **0.01** | | | | |

Abbreviations: PBVC, percentage brain volume chance; PVVC, percentage ventricle volume change; T0, baseline; T1, ~5-year follow-up; T2, ~13-year follow-up; S.E., standard error; df, degrees of freedom; Adj., adjusted; BMI, body mass index; ISI, interscan interval.

### Table S7| Summary of SIENAX total brain volume, antipsychotic class and nervous system drug linear mixed effect model.

| **Fixed Effects** | **Estimate** | **S.E.** | **t-value** | **p-values** |
|---|---|---|---|---|
| ATC-N | 8171.84 | 3223.77 | 2.54 | **0.01** |
| FGA | -26526.20 | 14016.71 | -1.89 | 0.06 |
| FGA + SGA | -25024.28 | 16636.06 | -1.50 | 0.14 |
| SGA | -31568.93 | 13184.63 | -2.39 | **0.02** |
| Time (continuous) | -5637.49 | 1023.25 | -5.51 | **3.55e-07** |
| Age at baseline | -3258.59 | 1055.51 | -3.09 | **3.05e-03** |
| Sex | 26195.52 | 19717.80 | 1.33 | 0.19 |
| BMI | -1217.91 | 1009.76 | -1.21 | 0.23 |
| Scanner | -21776.16 | 11862.18 | -1.84 | 0.07 |
| **Random Effects** | **Variance** | **Std. Dev.** | | |
| Participant (Intercept) | 3349084428.72 | 57871.28 | | |
| Residuals | 644876254.32 | 25394.41 | | |

Abbreviations: ATC-N, nervous system drugs; FGA, first-generation antipsychotics; SGA, second-generation antipsychotics; BMI, body mass index; S.E., standard error; Std. Dev., standard deviation.

### *Figures*

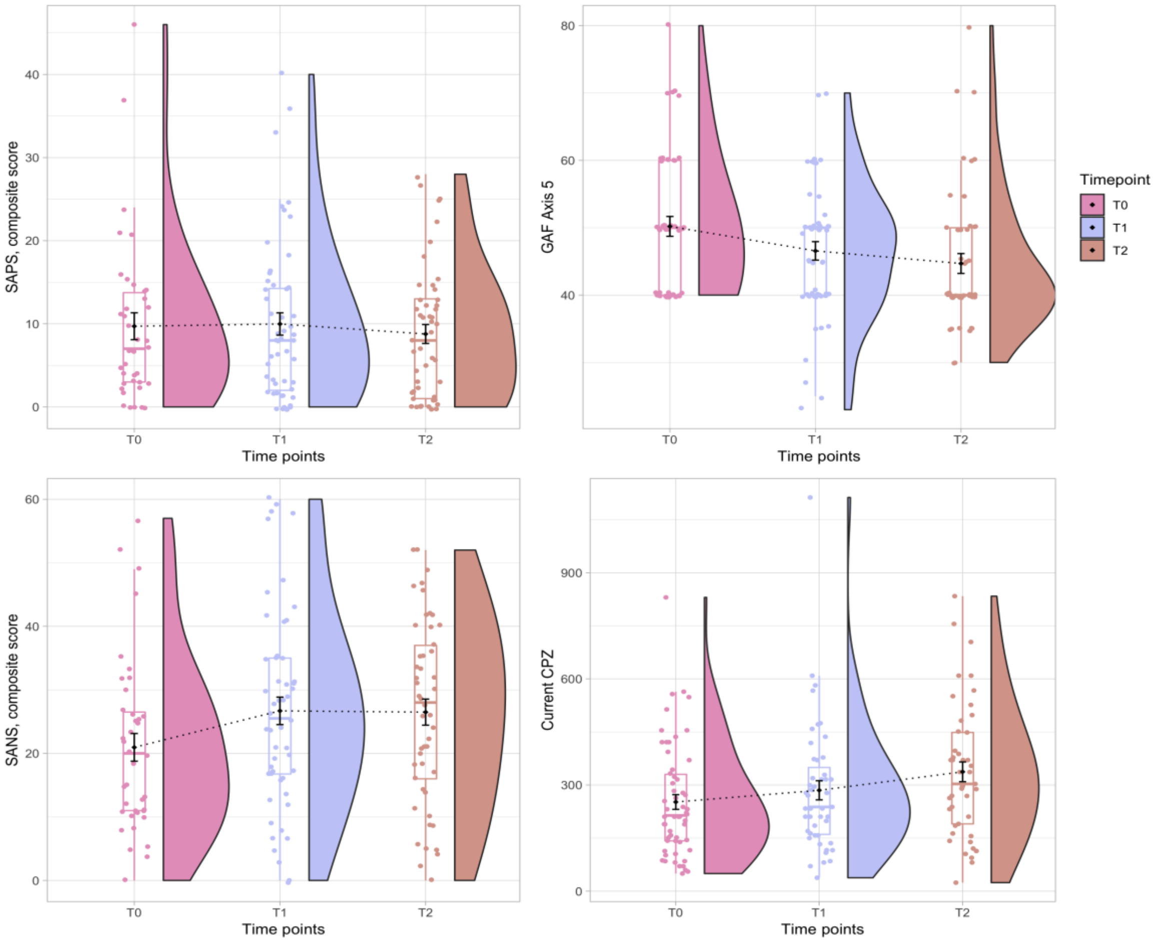


**Figure S1| Change in clinical variables over time in patients.** The data is displayed as a raincloud plot, which combines boxplots, raw data point and the distribution of data using split-half violins. Abbreviations: SAPS, scale for assessment of positive symptoms; GAF, global assessment of functioning; SANS, scale for assessment of negative symptoms; CPZ, chlorpromazine equivalent; T0, baseline; T1, ~5-year follow-up; T2, ~13-year follow-up.

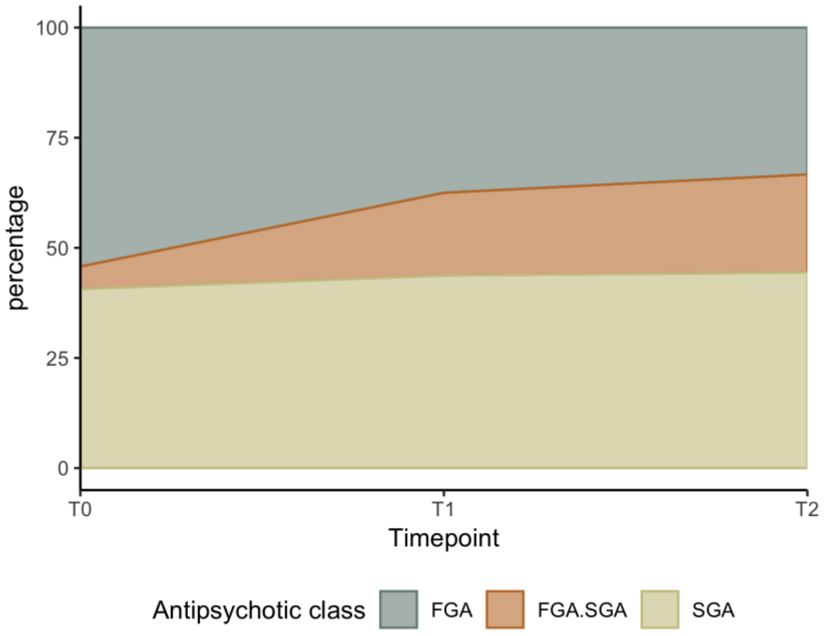


**Figure S2| Usage of first- (FGA) and/or second-generation antipsychotics (SGA) over time.** Area charts of percentages are displayed.

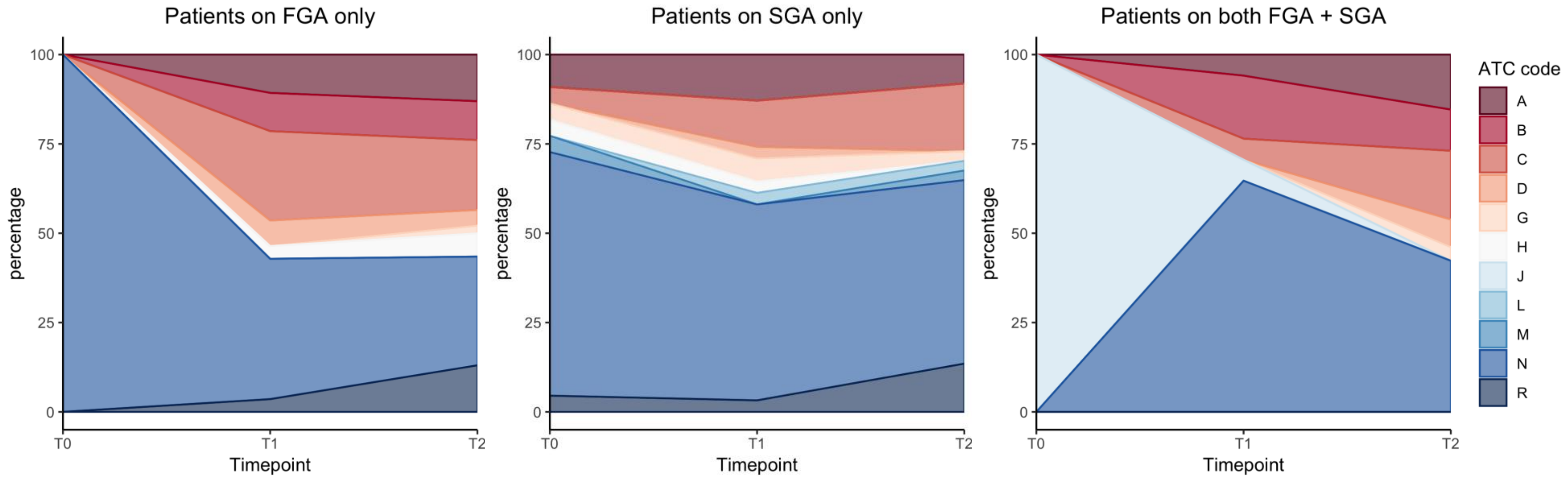

**Figure S3| Overall use of medication in patients, stratified by antipsychotic medication class.** Area chart of counts for each medication type used at each time point, stratified by antipsychotic medication class, based on the anatomical therapeutic chemical classification system, first-level (first panel).

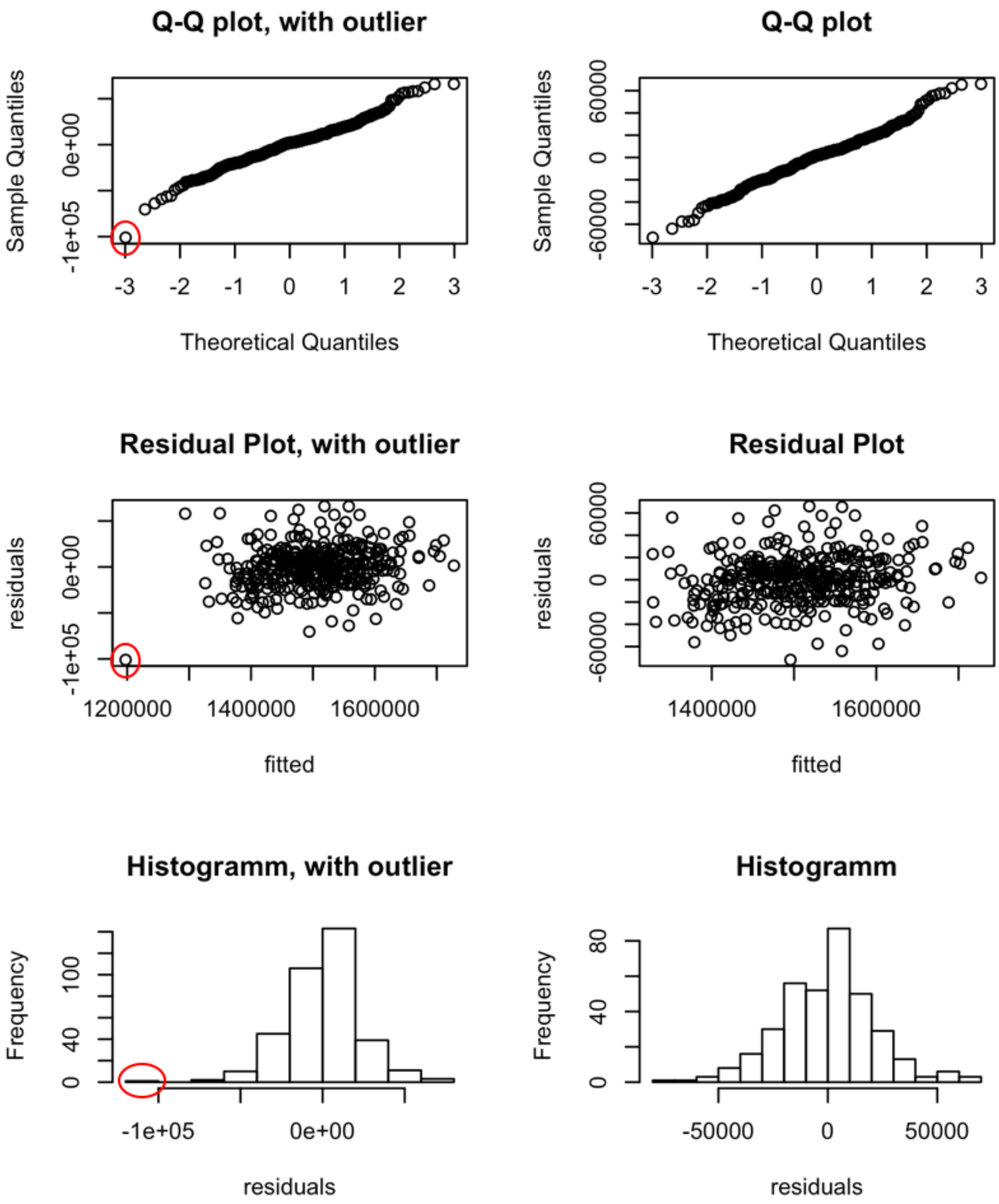


**Figure S4| Diagnostic plots of SIENAX total brain volume linear mixed effect model with (left panel) and without (right panel) outlier.** The identified influential outlier is marked with a red circle.

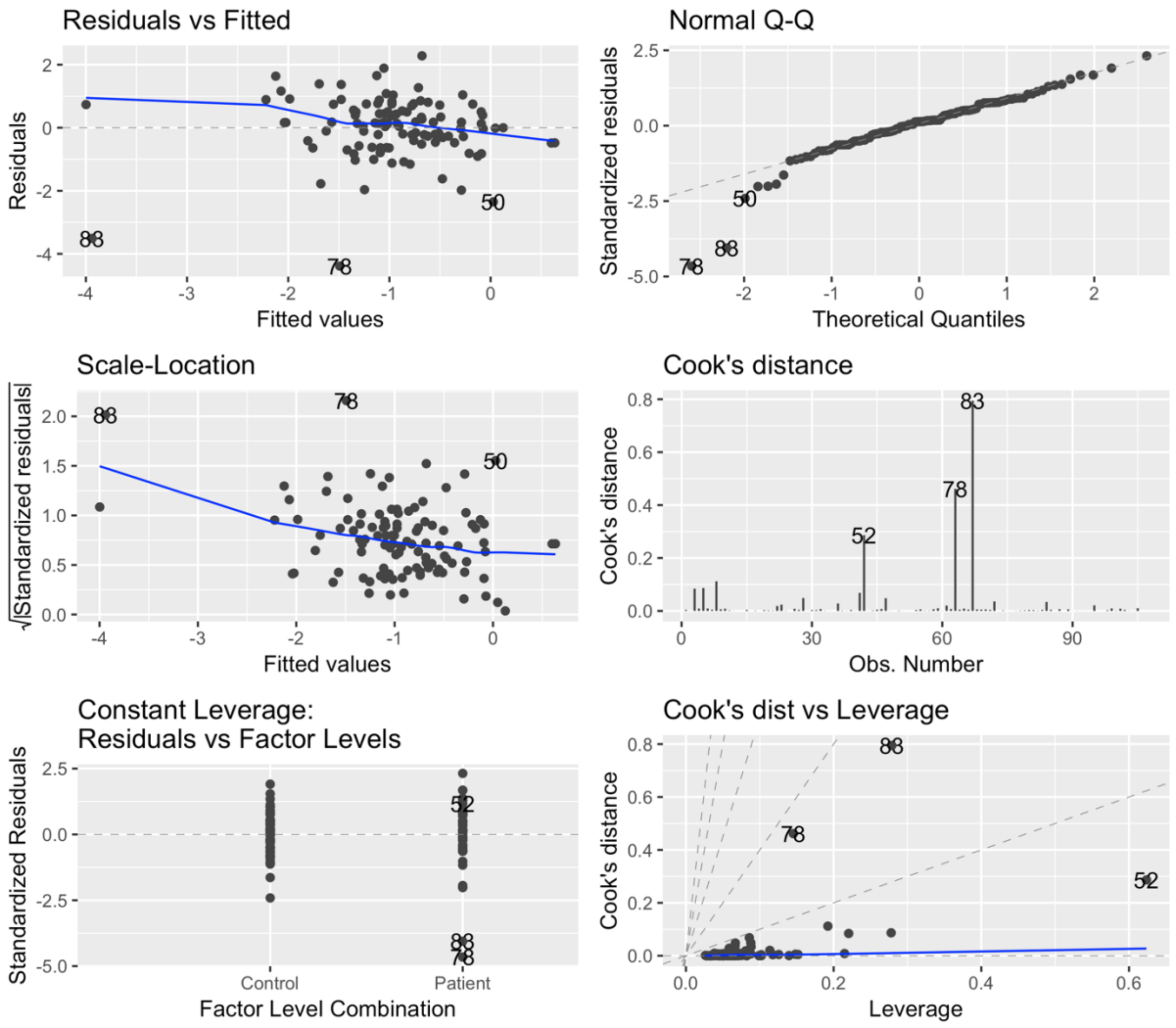


**Figure S5| Diagnostic plots of multiple linear regression model with percentage brain volume change between baseline and ~5-year follow-up as dependent variable.** The identified influential outliers are number 78 and 83.

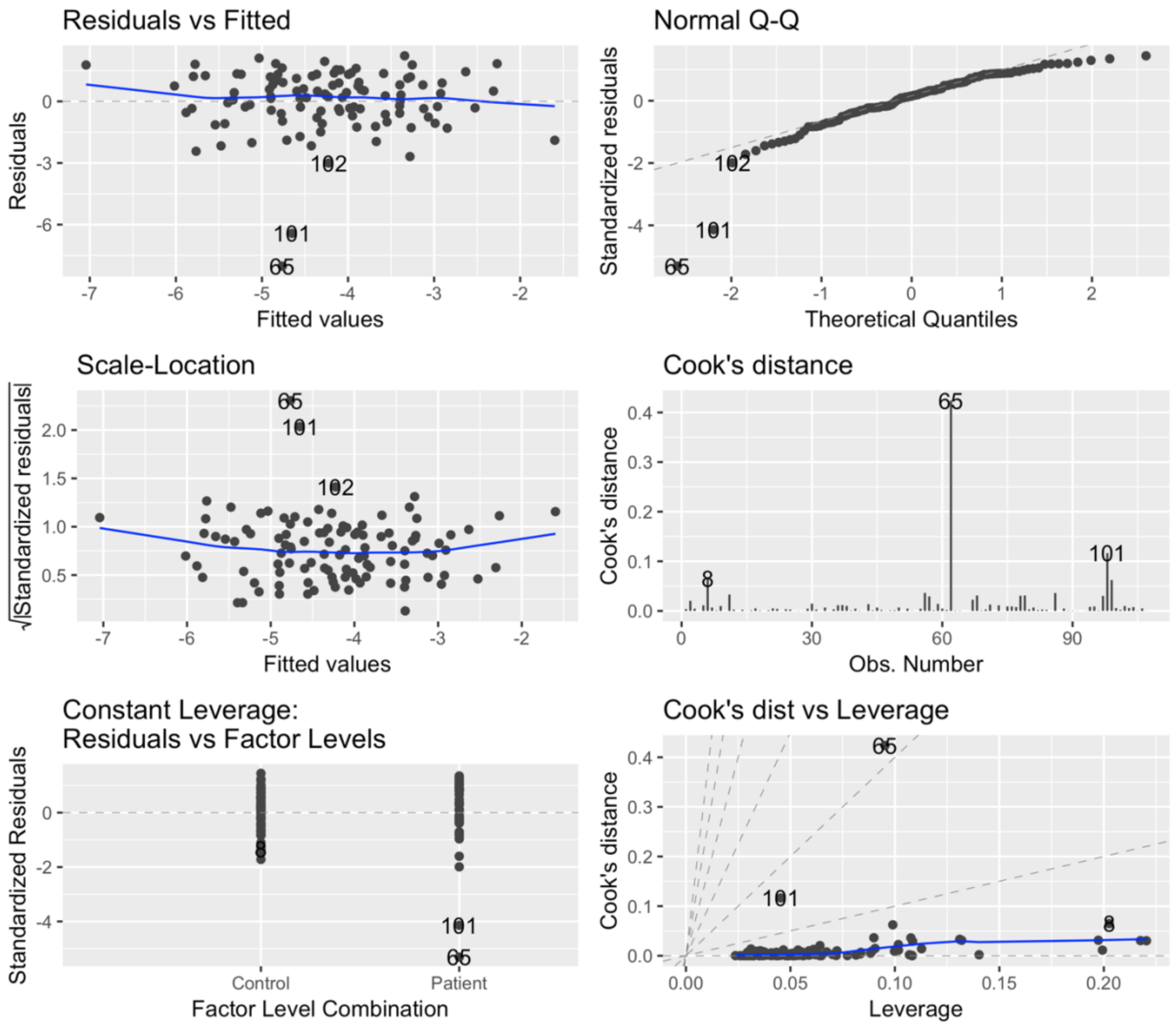


**Figure S6| Diagnostic plots of multiple linear regression model with percentage brain volume change between baseline and ~13-year follow-up as dependent variable.** The identified influential outlier is number 65.

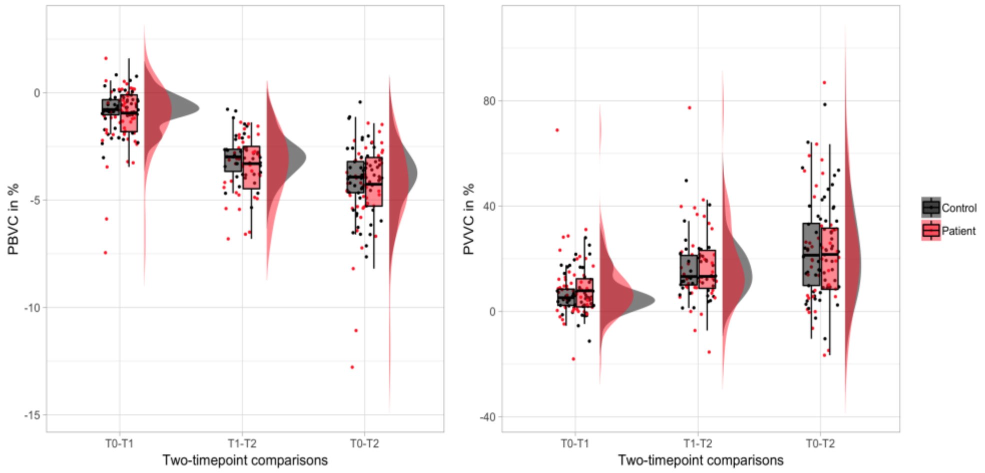


**Figure S7| Percentage brain (PBVC) and ventricle volume change (PVVC) over time.** The data is displayed for PBVC (left, including outlier) and PVVC (right) as a raincloud plot, which combines boxplots, raw data point and the distribution of data using split-half violins[48]. Patients are displayed in red and controls in black.

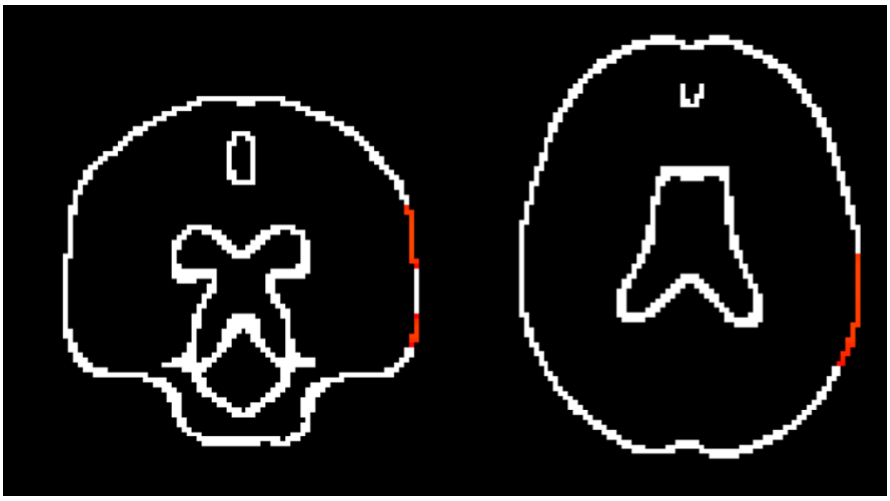

**Figure S8| Case-control differences in atrophic brain edge points for two timepoint comparison from baseline to ~13-year.** Significant brain edge points, which are atrophic in patients relative to controls, are overlaid on standard MNI152-space brain edge image. The analysis was co-varied for baseline age, time (continuous), sex and body mass index change.

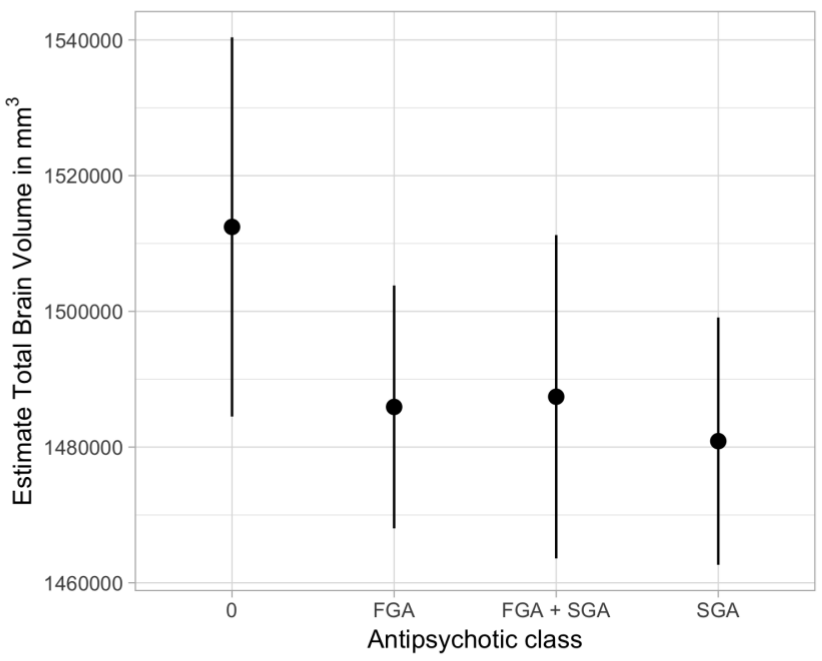


**Figure S9| Estimates of total brain volume for the interaction between antipsychotic class and non-antipsychotic nervous-system drugs.** Values are corrected for time (continuous), age at baseline, sex, body mass index and scanner. Estimates are displayed with upper and lower confidence intervals. 0 entails no medication at time points assessed.

### References


1. Nesvag R, Schaer M, Haukvik UK, et al. Reduced brain cortical folding in schizophrenia revealed in two independent samples. *Schizophr Res.* 2014;152(2-3):333-338.
2. Vares M, Ekholm A, Sedvall GC, Hall H, Jonsson EG. Characterization of patients with schizophrenia and related psychoses: evaluation of different diagnostic procedures. *Psychopathology.* 2006;39(6):286-295.
3. Smith SM, De Stefano N, Jenkinson M, Matthews PM. Normalized accurate measurement of longitudinal brain change. *J Comput Assist Tomo.* 2001;25(3):466-475.
4. Smith SM, Zhang Y, Jenkinson M, et al. Accurate, robust, and automated longitudinal and cross-sectional brain change analysis. *Neuroimage.* 2002;17(1):479-489.
5. Smith SM. Fast robust automated brain extraction. *Hum Brain Mapp.* 2002;17(3):143-155.
6. Jenkinson M, Bannister P, Brady M, Smith S. Improved optimization for the robust and accurate linear registration and motion correction of brain images. *Neuroimage.* 2002;17(2):825-841.
7. Jenkinson M, Smith S. A global optimisation method for robust affine registration of brain images. *Med Image Anal.* 2001;5(2):143-156.
8. Zhang Y, Brady M, Smith S. Segmentation of brain MR images through a hidden Markov random field model and the expectation-maximization algorithm. *IEEE Trans Med Imaging.* 2001;20(1):45-57.
9. Bartsch AJ, Homola G, Biller A, et al. Manifestations of early brain recovery associated with abstinence from alcoholism. *Brain.* 2007;130(Pt 1):36-47.
